# Chemical proteomics reveals post-translational remodeling of endosomal pathways during macrophage activation

**DOI:** 10.64898/2026.09.25.754469

**Authors:** Tiffany L. Zhang, Henry Sanford, Railia Gabbasova, Anne E. van Vlimmeren, Yannis Kalaidzidis, Hiroyuki Kojima, Charlotte R. Wayne, Jeetayu Biswas, Priyam Banerjee, Beatrice Zhang, Omar Abdel-Wahab, Marino Zerial, Ekaterina V. Vinogradova

## Abstract

Macrophages are central regulators of innate immunity, yet the post-translational mechanisms that enable rapid inflammatory responses remain incompletely understood. Here, we integrated transcriptomics, unenriched proteomics, phosphoproteomics, metabolomics, and cysteine reactivity profiling to comprehensively characterize primary human macrophages following Toll-like receptor and Stimulator of Interferon Genes (STING) activation. This approach revealed substantial discordance between transcript- and protein-level changes, as well as extensive post-translational remodeling. Differentially reactive cysteines were enriched in proteins involved in small-GTPase signaling, vesicular trafficking, and the endo-lysosomal system. Reactivity changes in two subunits of the endosomal mRNA-transport FERRY complex nominated it as a candidate target of inflammation-associated post-translational regulation. Biochemical and imaging analyses established FERRY complex assembly and early endosomal localization in primary human macrophages and identified DISC1 as a previously unrecognized FERRY-associated protein. This work provides a comprehensive resource of biochemical and post-translational remodeling during macrophage activation and implicates endosome-associated mRNA transport in innate immune regulation.

## INTRODUCTION

Macrophages coordinate innate immune responses by integrating environmental cues to regulate antigen presentation, cytokine production, and phagocytosis^1^. Toll-like receptors (TLRs)^2^ and the stimulator of interferon genes (STING) pathway^3^ are complementary innate immune sensors that detect diverse pathogen- and damage-associated molecular patterns. Activation of these pathways represents a central mechanism driving pro-inflammatory macrophage signaling. Dysregulation of TLR and STING signaling has been implicated in numerous human diseases^4, 5^. Autoimmune disorders, including systemic lupus erythematosus and rheumatoid arthritis^6^, are characterized by elevated levels of endogenous TLR ligands, whereas defects in downstream TLR signaling components have been linked to increased susceptibility to infection, attenuated innate immune responses, and impaired T cell activation^7^.

Despite the central role of these pathways in host defense and disease, their molecular characterization has relied heavily on immortalized cell lines and mouse models that fail to fully recapitulate the biology of primary human macrophages^8, 9, 10^. Furthermore, most studies have focused on transcriptomic profiling, even though mRNA levels often correlate poorly with protein abundance^11, 12, 13, 14^. Protein function is further regulated through post-translational mechanisms that play a role in their activity, localization, and molecular interactions. As a result, abundance-based proteomic analyses alone provide an incomplete picture of the biochemical changes that control macrophage activation.

To address this gap, we generated a comprehensive multi-omic atlas of TLR- and STING-activated primary human macrophages by integrating transcriptomics, abundance-based and functional proteomics, and metabolomics. Central to this analysis was tandem mass tag-enabled activity-based protein-profiling (TMT-ABPP)^15, 16^, which uses cysteine-reactive probes to detect structural and functional changes in protein states independently of protein abundance. Previous studies have demonstrated that this approach captures proteome remodeling associated with phosphorylation^17^, redox signaling during T cell activation^15^, metabolic dysfunction associated with T cell exhaustion^18^, and lysosomal trafficking in macrophages infected with *Mycobacterium tuberculosis*^19^. In dysfunctional T cells, this approach also identified broader protein networks sensitive to ATP availability^18^, highlighting the capacity of reactive cysteines to serve as sensitive reporters of cellular biochemical state beyond changes in protein abundance.

Here, we applied this integrated strategy to characterize diverse pro-inflammatory states induced by TLR and STING agonists in primary human macrophages. The resulting dataset revealed extensive proteome remodeling, including coordinated changes in small GTPase networks, vesicle trafficking machinery, and phosphoinositide signaling across the endo-lysosomal system. Among the state-dependent targets, we identified two subunits of FERRY, a recently described Rab5 effector complex that mediates endosomal mRNA transport and localized translation in neurons^20, 21^. We further demonstrated FERRY complex assembly and TLR4-dependent remodeling in macrophages, pointing to a previously unrecognized role of endosome-associated mRNA transport and localized translation in macrophage biology.

## RESULTS

### A scalable primary human macrophage platform enables multi-omic profiling of TLR and STING activation

We developed a robust and scalable *in vitro* differentiation platform to characterize molecular programs associated with TLR and STING signaling in primary human macrophages. Briefly, CD14^+^ monocytes isolated from peripheral blood mononuclear cells (PBMCs) of healthy donors were differentiated into resting (M0) macrophages over six days in the presence of macrophage colony stimulating factor (mCSF), followed by stimulation with TLR ligands or a STING agonist for 24 hours before downstream analyses (Fig. 1a). This platform enables generation of primary human macrophages, supporting integrated multi-omic profiling of macrophage activation, including transcriptomics, abundance-based proteomics, cysteine reactivity profiling, phosphoproteomics, and metabolomics.

**Fig. 1.**
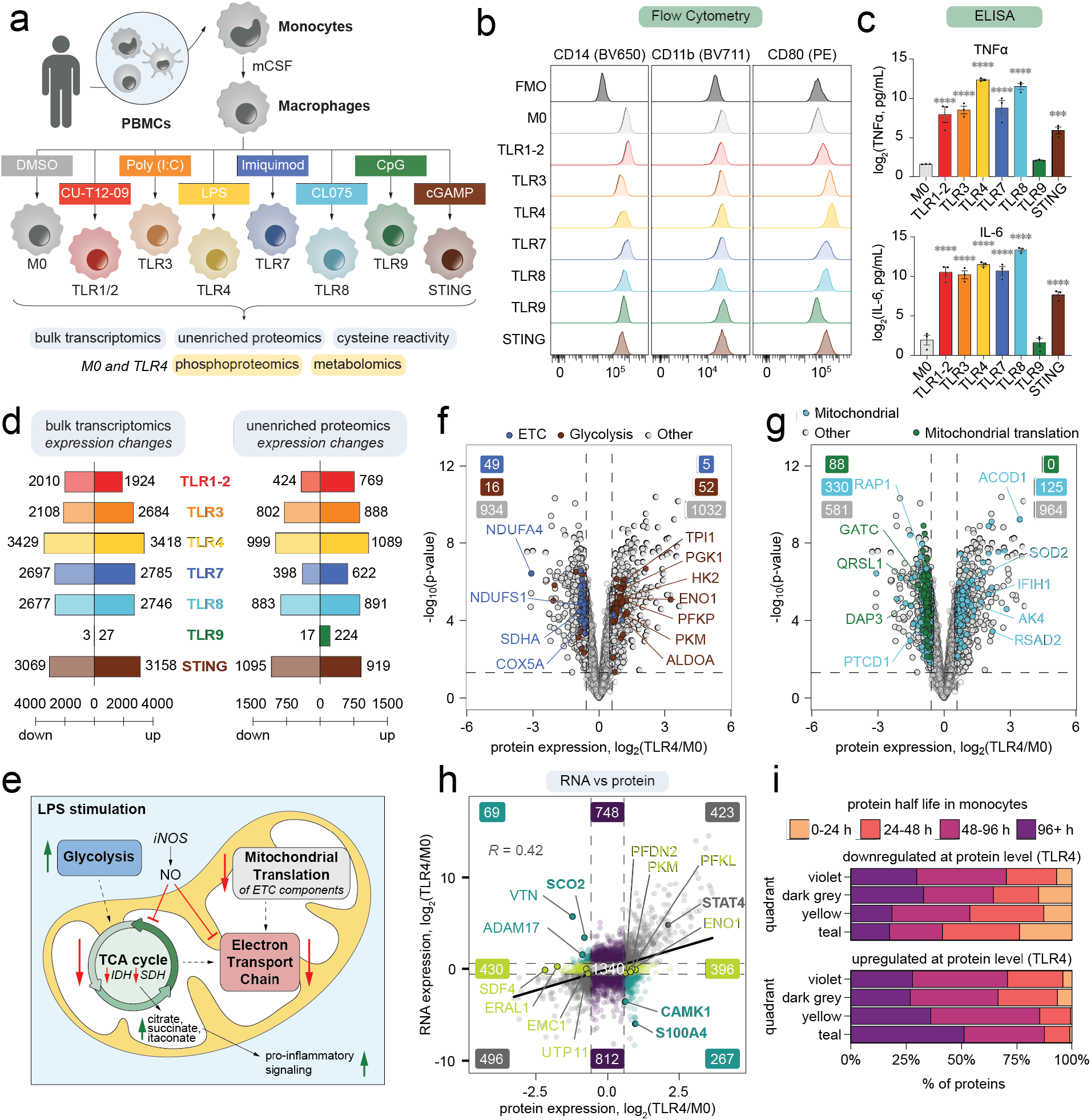
Transcriptomic and proteomic profiling of TLR- and STING-stimulated macrophages. **a**, Schematic of macrophage stimulation conditions and multi-omic platforms used to profile activated macrophages. **b**, Representative flow cytometry histograms for cell-surface expression of CD14, CD11b, and CD80 across macrophage activation states. **c**, Secreted TNF and IL-6 levels measured by ELISA in culture supernatants following 24 h of macrophage stimulation. **d**, Numbers of upregulated and downregulated transcripts and proteins identified by bulk RNA-seq (left; p < 0.05, FC > 2, n = 3 donors) and abundance-based proteomics (right; p < 0.05, FC > 1.5, n = 3 donors), respectively. **e**, Schematic illustrating metabolic rewiring in pro-inflammatory macrophages. **f**,**g**, Volcano plots showing log_2_(fold change) of proteins between TLR4-stimulated and unstimulated M0 cells. The horizontal dashed line marks p = 0.05 and the vertical dashed lines mark a 1.5-fold change. Proteins associated with glycolysis (Hallmark) and the electron transport chain (ETC; Gene Ontology) are highlighted according to their respective categories in (**f**), and proteins associated with mitochondria and mitochondrial translation are highlighted in (**g**). **h**, Scatter plot comparing log_2_(fold change) in protein abundance (x-axis) and transcript abundance (y-axis) between TLR4-stimulated and unstimulated M0 macrophages. Proteins are colored teal, yellow, purple or grey according to the concordance between protein- and transcript-level changes. Pearson’s correlation coefficient and regression line are shown. Only genes quantified in both proteomic and transcriptomic datasets are included. Transcriptomic and proteomic data are from n = 3 donors. **i**, Reference protein half-lives in monocytes^24^ for upregulated and downregulated proteins, depending on their correlation with transcript-level changes. Half-life values represent the mean of two biological replicates, and only proteins with reported half-life data are included.

We validated this platform by assessing macrophage activation using flow cytometry analysis of cell-surface markers and quantification of inflammatory cytokine secretion by ELISA (Fig. 1b,c and Extended Data Fig. 1a). Stimulation of multiple TLR pathways resulted in reduced surface levels of CD14, consistent with the reported endocytosis of this receptor following activation^22^. CD11b remained relatively unchanged, while the costimulatory receptor CD80 was upregulated in most stimulation conditions (Fig. 1b and Extended Data Fig. 1a). In parallel, we observed elevated secretion of pro-inflammatory cytokines IL-6 and TNF, canonical downstream effectors of TLR signaling in macrophages^23^ (Fig. 1c). These results confirmed robust activation of primary human macrophages across the stimulation conditions, with the notable exception of TLR9 stimulation, which did not show altered surface marker expression or cytokine production relative to M0 cells.

### TLR and STING activation drive extensive inflammatory and metabolic remodeling in primary human macrophages

To comprehensively characterize molecular programs associated with macrophage activation, we profiled transcriptional and proteomic changes using RNA-sequencing and abundance-based proteomics with tandem mass tag-enabled sample multiplexing (TMT-exp; Fig. 1d and Extended Data Fig. 1b,c; Data S1-1, S2-1, S2-2). Transcriptional responses were generally more pronounced than corresponding changes in protein abundance, underscoring the contribution of post-transcriptional mechanisms to macrophage activation and highlighting the importance of protein-level analyses for capturing functionally relevant transitions in macrophage states. Consistent with our immune profiling experiments, TLR9 agonism produced minimal transcriptional and protein level changes, likely reflecting low TLR9 expression in mCSF-differentiated macrophages. Consistent with this, TLR9 itself was not quantified in our RNA-sequencing and abundance-based proteomics experiments, and this condition was therefore excluded from subsequent analyses.

Principal component analysis of both transcriptomic and proteomic profiles revealed clear separation of TLR- and STING-stimulated macrophages from resting M0 cells (Extended Data Fig. 1b,c). Within the TLR conditions, TLR3- and TLR4-stimulated macrophages clustered separately from other TLR-stimulated conditions, consistent with their ability to engage TRIF-dependent signaling and induce type I interferon responses, in contrast to the predominantly MyD88-dependent pathways used by other TLRs^23^. STING activation, which also promotes type I interferon production through a mechanistically distinct pathway, generated a discrete transcriptional and proteomic signature, clustering separately from both TLR-stimulated and resting macrophages, highlighting the diversity of pro-inflammatory states captured within this dataset.

Gene set enrichment analysis (GSEA) of proteins contributing to the principal component 1 (PC1) loadings revealed upregulation of pro-inflammatory and antiviral programs and downregulation of aerobic metabolic pathways (Extended Data Fig. 1d; Data S2-3). Principal component 2 (PC2) distinguished STING activation from other TLR-stimulated conditions, highlighting the unique molecular features of the STING response. Consistent with these findings, GSEA demonstrated broad activation of TNF signaling across all inflammatory states, while interferon-responsive programs were preferentially enriched following STING, TLR3, and TLR4 stimulation (Extended Data Fig. 1e,f).

Upon activation, macrophages undergo metabolic reprogramming characterized by a shift from oxidative phosphorylation (OXPHOS) to aerobic glycolysis^25^. This transition supports rapid ATP generation and is accompanied by remodeling of the tricarboxylic acid (TCA) cycle, leading to accumulation of immunoregulatory metabolites, including citrate^26^, succinate^27, 28^, and itaconate^29, 30^ (Fig. 1e). In parallel, increased glycolytic flux and diversion of glucose intermediates into the pentose phosphate pathway promote NADPH production and reactive oxygen species (ROS) generation^25^. Consistent with this, we observed increased expression of glycolytic enzymes across all stimulation conditions, together with reduced abundance of multiple electron transport chain (ETC) components and ATP synthase subunits (Fig. 1f,g, Extended Data Fig. 1g-i).

Notably, macrophage activation was associated with broad suppression of mitochondrial translation, in contrast to the more heterogeneous regulation of other mitochondrial proteins (Fig. 1g and Extended Data Fig. 1i). Downregulated proteins included mitochondrial ribosome-associated GTPases (ERAL1), components of mitochondrial tRNA processing complexes (PTCD1, QRSL1, GATC), and numerous mitochondrial ribosomal subunits. Since mitochondria encode and synthesize key subunits of the ETC, coordinated suppression of ETC components and mitochondrial translation may enable macrophages to rapidly downregulate OXPHOS during inflammation. These findings underscore the extensive metabolic remodeling that is required during TLR- and STING-mediated macrophage activation and the establishment of distinct inflammatory cell states.

### Post-transcriptional regulation contributes to proteome remodeling during macrophage activation

We next compared transcript- and protein-level changes to identify proteins subject to post-transcriptional regulation and define factors contributing to this divergence (Fig. 1h, Extended Data Fig. 2a; Data S2-2). In TLR4-stimulated macrophages, we observed only moderate correlation between mRNA and protein abundance (R = 0.42; Fig. 1h). Similar correlations were found for other stimulation conditions in macrophages (Extended Data Fig. 2a), in line with our observations in activated and exhausted T cells^15, 18^. This analysis identified proteins whose abundance closely tracked transcript levels (grey), as well as proteins regulated predominantly at either the transcript (purple) or protein (yellow) level, in addition to a subset of proteins displaying anti-correlated expression patterns (teal) (Fig. 1h).

Gene ontology analysis of proteins in the teal (anti-correlated) and yellow (regulated at the protein-level) categories revealed enrichment of metabolic, signaling, and mitochondrial processes, including glycolysis (PKM, PFKL, ENO1) and mitochondrial translation (mitochondrial ribosomes and aminoacyl-tRNA synthetases) (Fig. 1h, Extended Data Fig. 2b; Data S2-4). The post-transcriptional regulation of these pathways may enable rapid adaptation to inflammatory and metabolic stress during macrophage activation. We validated representative targets by qPCR and Western blot analysis, including SCO2, CAMK1, and S100A4 (teal), as well as STAT4 (grey) (Extended Data Fig. 2c-e).

Our findings indicate that transcriptional regulation accounts for only a subset of the proteomic remodeling that accompanies macrophage activation, suggesting additional layers of post-translational regulation that shape inflammatory protein landscapes. To test whether differences in protein turnover also contribute to the observed protein-transcript discordance, we integrated published protein half-life measurements from primary human monocytes^24^. Proteins selectively downregulated following TLR4 stimulation at the protein level (teal) were enriched for short-lived species, with relatively few exhibiting half-lives greater than 48 h (Fig. 1i, top). Conversely, proteins selectively upregulated at the protein level tended to be longer lived, with fewer than 20% displaying half-lives below 48 h (Fig. 1i, bottom). This trend was conserved across all stimulation conditions (Extended Data Fig. 2f), suggesting that protein turnover may serve as an additional layer of regulation of protein abundance and ultimately, macrophage activation states.

### Cysteine reactivity profiling reveals extensive biochemical and post-translational remodeling during macrophage activation

To capture additional layers of post-translational regulation during macrophage activation, we applied TMT-ABPP^15, 16^, a multiplexed chemical proteomic platform that uses cysteine reactivity toward electrophilic probes as a readout of biochemical changes in distinct cellular states (Fig. 2a; Data S2-5). The broadly reactive iodoacetamide-desthiobiotin (IA-DTB) probe covalently labels a subset of cysteine residues, which can then be selectively enriched for downstream proteomic analysis. Probe reactivity is sensitive to post-translational modifications (PTMs)^17^, changes in redox state^15^, protein-metabolite and protein-protein interactions (PPIs)^18^, while enrichment of cysteine-reactive peptides can also be indirectly affected through modifications of adjacent amino acids within the same tryptic peptide^17^. Thus, TMT-ABPP provides an unbiased and orthogonal strategy for identifying different forms of biochemical and post-translational regulation that shape macrophage activation states.

**Fig. 2.**
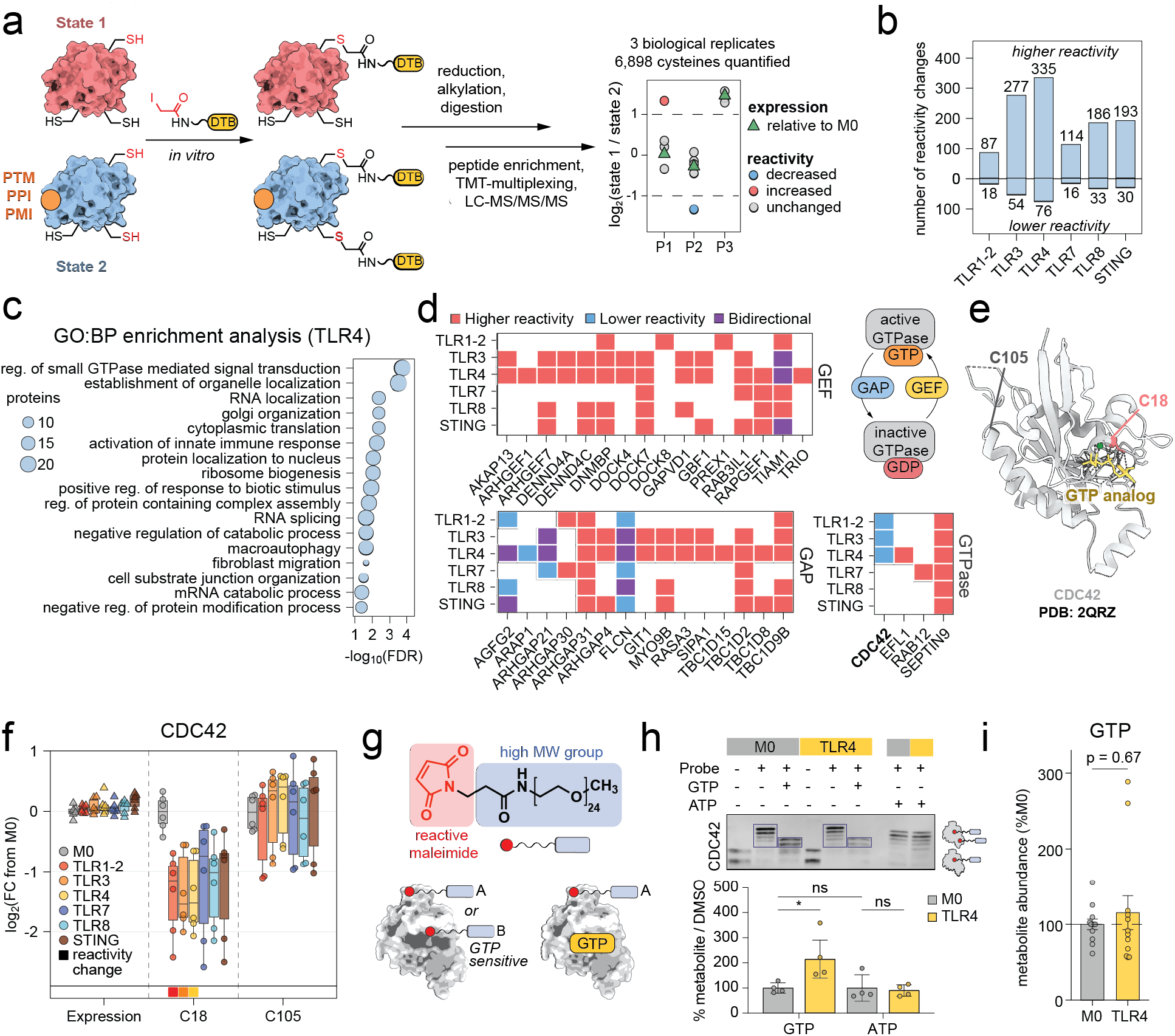
Cysteine reactivity profiling of TLR- and STING-stimulated macrophages reveals remodeling of GTPases and their regulators. **a**, Schematic of cysteine reactivity profiling and representative plot illustrating expression and reactivity changes. PTM, post-translational modification; PPI, protein-protein interaction; PMI, protein-metabolite interaction. **b**, Numbers of cysteines with reactivity changes across macrophage stimulation conditions relative to M0 cells (n = 3 donors). Differential reactivity was determined relative to changes in protein abundance measured by TMT-exp, when available, or relative to the median cysteine reactivity change for the corresponding protein. **c**, GO:BP analysis (Fisher’s exact test) of proteins with cysteine reactivity changes in TLR4-stimulated macrophages relative to M0 cells. Terms with FDR < 0.01 and protein number > 5 are shown. Reg., regulation. **d**, Heatmaps showing cysteine reactivity changes across macrophage stimulation conditions for proteins grouped by functional class: guanine nucleotide exchange factors (GEFs), GTPase-activating proteins (GAPs), and GTPases. Colors indicate the direction of reactivity changes relative to protein expression or median cysteine reactivity. Blank cells indicate no reactivity change. **e**, Crystal structure of CDC42 bound to a GTP analog (yellow; PDB 2QRZ^31^), with the differentially reactive cysteine C18 highlighted in red and the non-reactive cysteine C105 in grey. **f**, Box plots showing changes in CDC42 expression (triangles) and cysteine reactivity (dots) across macrophage stimulation conditions compared to M0 cells. Boxes indicate the 25th to 75th percentiles, and whiskers extend to 1.5x the interquartile range. Boxes on the bottom margin indicate cysteines classified as differentially reactive. **g**, Structure of the high-molecular-weight, cysteine-reactive maleimide probe used for Western blot - Activity-based protein profiling (WB-ABPP) experiments (top), and schematic showing potential covalent adducts formed following probe treatment, including modification of GTP-sensitive cysteines (bottom). **h**, Representative CDC42 Western blot of lysates from M0 or TLR4-stimulated macrophages treated with GTP or ATP before labeling with the maleimide probe (top), with quantification of CDC42 bands shown below. **i**, GTP abundance in M0 and TLR4-stimulated macrophages. Data in (**h**) and (**i**) represent the average ± SEM of n = 4 donors, and statistical comparison is by a two tailed t-test with p-values shown (ns, p > 0.05; *, p < 0.05).

We compared cysteine reactivity across resting (M0), TLR-, and STING-stimulated macrophages using a 16-plex TMT-ABPP workflow (Fig. 2a,b; Data S2-5). To identify robust reactivity changes independent of protein abundance changes, we applied a stringent set of filters, restricting analysis to proteins with at least two quantified cysteine-containing peptides and reactivity changes exceeding twofold relative to the corresponding change in protein abundance (See Methods). Across all stimulation conditions, we identified 495 reactivity changes, with TLR3 and TLR4 stimulation producing the largest number of changes (Fig. 2b). Consistent with distinct cell-type-specific signaling programs, overlap in cysteine reactivity changes between macrophages and activated T cells^15^ was limited, likely reflecting differences in both protein expression and the underlying biology of these immune cell populations (Extended Data Fig. 3a,b).

### Chemoproteomic profiling reveals dynamic remodeling of GTPase signaling in activated macrophages

TLR4 stimulation produced the largest number of cysteine reactivity changes (Fig. 2b) and was therefore chosen for subsequent analyses. GO biological process (GO:BP) analysis of proteins with reactivity changes following TLR4 stimulation identified GTPase-mediated signal transduction as the most enriched term (Fig. 2c; Data S2-4). GTPases are central regulators of macrophage function, controlling inflammatory signaling, cytoskeletal dynamics, phagocytosis, and intracellular trafficking.^32, 33, 34, 35^ Closer inspection revealed >30 changes in GTPases, as well as in guanine nucleotide exchange factors (GEFs) and GTPase activating proteins (GAPs), which dynamically control GTPase activity (Fig. 2d).

GAPs and GEFs are extensively regulated by PTMs, PPIs, and second messenger binding. These regulatory events are frequently coupled to conformational changes^36, 37^ that can alter cysteine accessibility or environment, consistent with the reactivity changes observed in our study. Several of these mapped to domains with established functional roles, including PIP_3_-binding domains within the ArfGAP ARAP1^38^ and the RacGEF TIAM1^39^, with PIP_3_ binding known to promote their catalytic activity (Fig. 2d). Other changes, including those in RabGEFs RAB3IL1 and DENND4C and the ArfGAP GIT1, occur in disordered regions adjacent to known phosphorylation sites, suggesting potential regulation through phosphorylation.

We also identified reactivity changes within the nucleotide-binding pockets of several GTPases, including CDC42 (Fig. 2e,f) and Rab12 (Extended Data Fig. 3c,d), suggesting that these changes may reflect GTP or GDP occupancy. CDC42 is a Rho family GTPase that regulates endocytosis, cytoskeletal remodeling and other cellular processes^40^. To test whether CDC42 cysteine reactivity is sensitive to guanine nucleotide binding, we performed *in vitro* GTP-addback experiments in lysates from M0 and TLR4-stimulated macrophages using a high molecular weight cysteine-reactive maleimide probe and monitored probe-induced band shifts by Western blot analysis. Probe labeling produced an upward shift in CDC42 migration, which was selectively blocked by GTP, but not ATP (Fig. 2g,h).

Notably, the GTP-sensitive band shift was more pronounced in M0 lysates compared to TLR4-stimulated lysates (Fig. 2g,h), consistent with a stimulation-dependent difference in CDC42 occupancy or regulatory state. We also observed reactivity changes in multiple CDC42 regulators, including ARHGEF7 and DOCK4/7, as well as in the CDC42 effector TNK2 (Extended Data Fig. 3e), suggesting remodeling of this pathway following TLR4 stimulation. Complementary metabolomic profiling of polar metabolites (Fig. 2i, Extended Data Fig. 3f, Data S3-1) revealed no change in intracellular GTP levels following TLR4 stimulation. This data suggests that the altered cysteine reactivity is unlikely to result from greater GTP availability and may instead reflect regulatory changes in nucleotide loading, conformation, or protein interactions of CDC42.

### Reactivity profiling reveals remodeling of vesicular trafficking and autophagy

GO:BP analysis revealed enrichment of proteins with reactivity changes in vesicular trafficking pathways in TLR4-stimulated macrophages (Fig. 3a; Data S2-4), including Golgi organization and macroautophagy. GO cellular component analysis (GO:CC) localized many of these changes to defined membrane-associated compartments, including early endosomes and autophagosomes (Fig. 3b). The proteins with reactivity changes span diverse roles in vesicle trafficking, including anterograde and retrograde transport, Golgi organization, vesicle tethering and fusion, and cytoskeletal regulation (Fig. 3c,d).

**Fig. 3.**
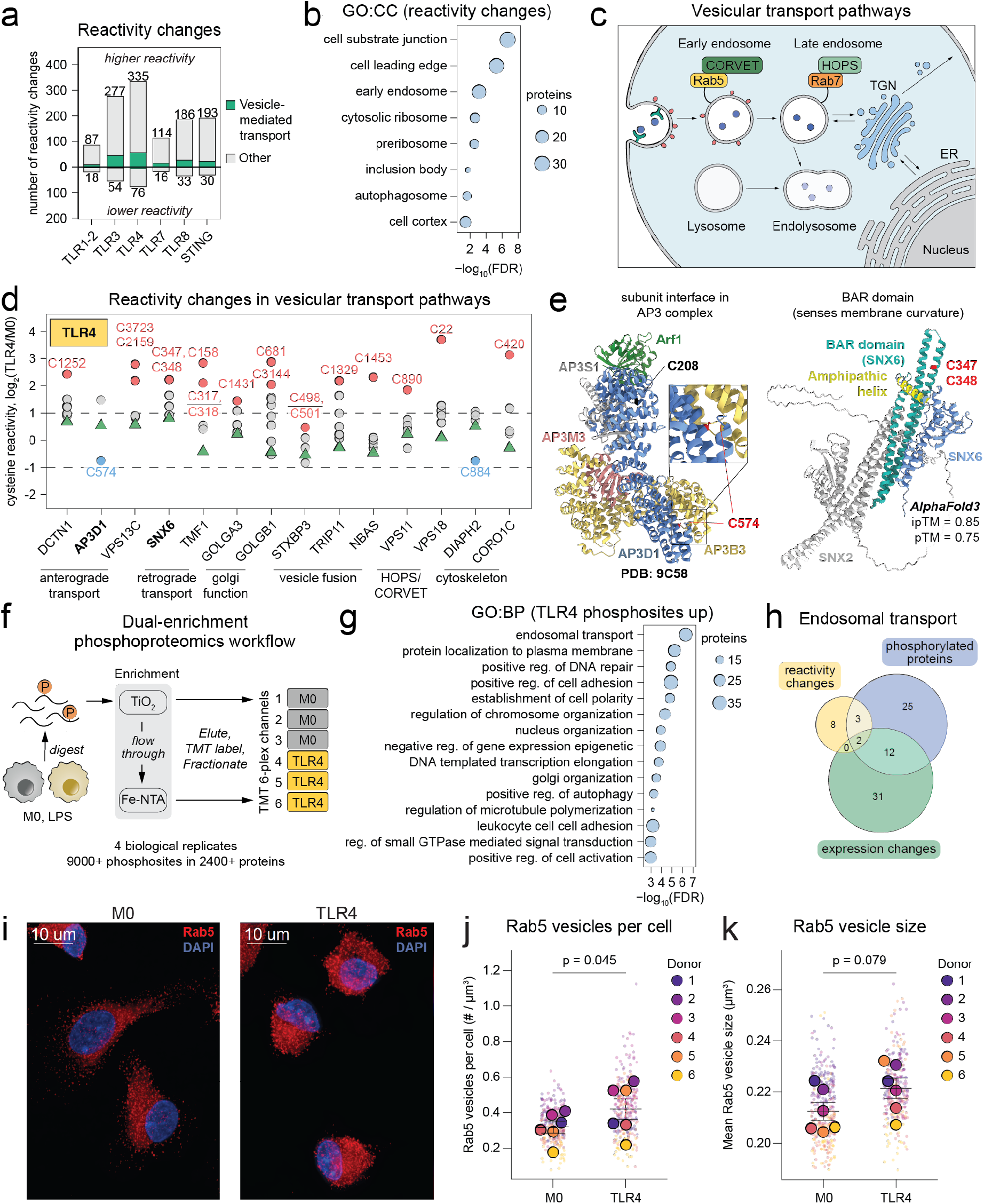
Cysteine reactivity and phosphoproteomic profiling reveal activation of vesicular trafficking pathways. **a**, Numbers of cysteines with reactivity changes across macrophage stimulation conditions relative to M0 cells (n = 3 donors). Proteins associated with vesicle-mediated transport based on Gene Ontology annotation are labeled in green. **b**, Gene ontology cellular component (GO:CC) enrichment analysis of proteins with cysteine reactivity changes in TLR4-stimulated macrophages relative to M0 cells. Terms with FDR < 0.01 and protein number > 5 are shown. Reg., regulation. **c**, Schematic showing vesicular trafficking pathways highlighting relevant organelles and select compartment-specific markers. **d**, Log_2_(fold change) in protein expression and cysteine reactivity for selected proteins in vesicular transport pathways in TLR4-stimulated macrophages relative to M0 cells. Protein expression changes are shown as green triangles and cysteine reactivity changes as dots. Cysteine reactivity and protein expression data are from n = 3 donors. **e**, Left, Cryo-EM structure of the AP3 complex containing AP3D1 (PDB 9C58^43^), with the differentially reactive cysteine C574 highlighted in red. Right, AlphaFold 3-predicted structure of SNX2-SNX6 heterodimer^44^, with C347 and C348, which are contained within the same differentially reactive tryptic peptide in SNX6, highlighted in red. **f**, Schematic of the dual-enrichment phosphoproteomics workflow that uses TiO_2_ and Fe-NTA enrichment to achieve in-depth phosphopeptide coverage. **g**, GO:BP enrichment analysis of proteins with phosphosites with significantly increased phosphorylation in TLR4-stimulated cells compared to M0 (FC > 2, p < 0.05). Terms with FDR < 0.01 and protein number > 5 are shown. **h**, Venn diagram of proteins associated with endosomal transport showing changes in cysteine reactivity (FC > 2 relative to expression value), expression (FC > 1.5, p < 0.05), or phosphorylation (FC > 2, p < 0.05) in TLR4-stimulated macrophages compared to M0 cells. **i**, Representative confocal microscopy images of Rab5 vesicles (red) with DAPI-stained nuclei (blue) in unstimulated M0 and TLR4-stimulated macrophages (Scale bar: 10 µm, zoomed: 5 µm). **j**, Quantification of number of vesicles per cell, normalized to total cell volume. Dots represent individual cells and are colored by donor. The average across cells for each donor is represented by a larger dot. Lines represent mean and SEM for n = 6 donors. Significance was determined by a two-tailed paired t-test on the donor means. **k**, Quantification of mean Rab5 vesicle size per cell. Dots represent individual cells and are colored by donor. The average across cells for each donor is represented by a larger dot. Lines represent mean and SEM for n = 6 donors. Significance was determined by a two-tailed paired t-test on the donor means.

Several reactivity changes mapped to functionally important structural domains, including AP3D1_C574, located at the interface of AP3B3 within the AP3 adaptor complex^41^, and C347/C348 in the BAR domain of SNX6^42^, a component of the retromer complex, which heteromerizes with SNX1 or SNX2 (Fig. 3d,e). We also identified reactivity changes in multiple core regulators of autophagy, including ATG9A, ATG7, and UVRAG, consistent with the established role of TLR signaling in autophagy^18^ (Extended Data Fig. 4a,b).

Membrane trafficking is coordinated by phosphatidylinositol lipids, which define organelle identity and direct protein recruitment through the concerted actions of lipid kinases and phosphatases^45^ (Extended Data Fig. 4c). Consistent with extensive remodeling of intracellular membrane compartments following TLR activation, we observed multiple reactivity changes in phosphatidylinositol regulators, including decreased reactivity of C312 in PIP5K1A following TLR3 and TLR4 stimulation (Extended Data Fig. 4d). This lipid kinase generates PI(4,5)P_2_ at the plasma membrane and nucleus and requires membrane recruitment and dimerization for functional activation^46, 47^. The differentially reactive cysteine C312 lies at the protein homodimer interface^46^ (Extended Data Fig. 4d,e), which is consistent with reported activation of this kinase under TLR4 stimulation conditions to modulate cytoskeletal activity at phagocytic cups^48^.

### Phosphoproteomic analysis provides complementary information on post-translational regulation in activated macrophages

Considering vesicular trafficking is a post-translationally regulated process^49, 50, 51^, we complemented our reactivity profiling with phosphoproteomics analysis of M0 and TLR4-stimulated macrophages (Fig. 3f; Data S2-6). Using a dual phosphopeptide enrichment workflow with TiO_2_ and Fe-NTA, we quantified more than 9,000 phosphosites across 2,498 proteins (Fig. 3f and Extended Data Fig. 5a-c). GO:BP analysis of phosphosites with altered abundance following TLR4 stimulation revealed significant enrichment of pathways involved in endosomal transport (Fig. 3g, Extended Data Fig. 5d-f, Data S2-4), GTPase-mediated signal transduction (Extended Data Fig. 5g,h), and vesicle-mediated transport (Extended Data Fig. 5i-k), among others. Notably, the proteins contributing to these pathways in the phosphoproteomic dataset were largely distinct from those identified by expression-based proteomics (TMT-exp) or cysteine reactivity profiling (TMT-ABPP) (Fig. 3h, Extended Data Fig. 5e-k), highlighting how these orthogonal approaches reveal complementary information on post-translational regulation of these pathways. Finally, immunofluorescence (IF) analysis of the subcellular localization and distribution of the early endosomal marker Rab5 confirmed increased abundance of early endosomes following TLR4 stimulation (Fig. 3i-k).

### Chemoproteomic profiling identifies FERRY complex as a post-translationally regulated target in activated macrophages

To prioritize cysteine reactivity changes most likely to reflect biologically relevant regulatory events, we examined the structural and evolutionary properties of differentially reactive cysteines (Data S2-5). Relative to all quantified cysteines, differentially reactive cysteines were enriched in solvent-accessible and intrinsically disordered regions predicted by AlphaFold2^52^ (Fig. 4a,b, Extended Data Fig. 6a,b) with a smaller fraction of these sites predicted to be pathogenic by AlphaMissense^53^. Most of the cysteines were also highly conserved across species in both reactive and nonreactive groups^54^. Notably, nearly half of the proteins with differentially reactive cysteines were annotated as components of known protein complexes (CORUM^55, 56^ and Complex Portal^57, 58^; Extended Data Fig. 6a), suggesting that many of the observed reactivity changes may reflect state-dependent structural rearrangements and remodeling of higher-order protein assemblies. We therefore prioritized reactive cysteines that were highly conserved^54^, predicted to be pathogenic by AlphaMissense, located proximal to known disease variants (ClinVar^59^), and mapped to proteins within known complexes^55, 56, 57, 58^ (Fig. 4a).

**Fig. 4.**
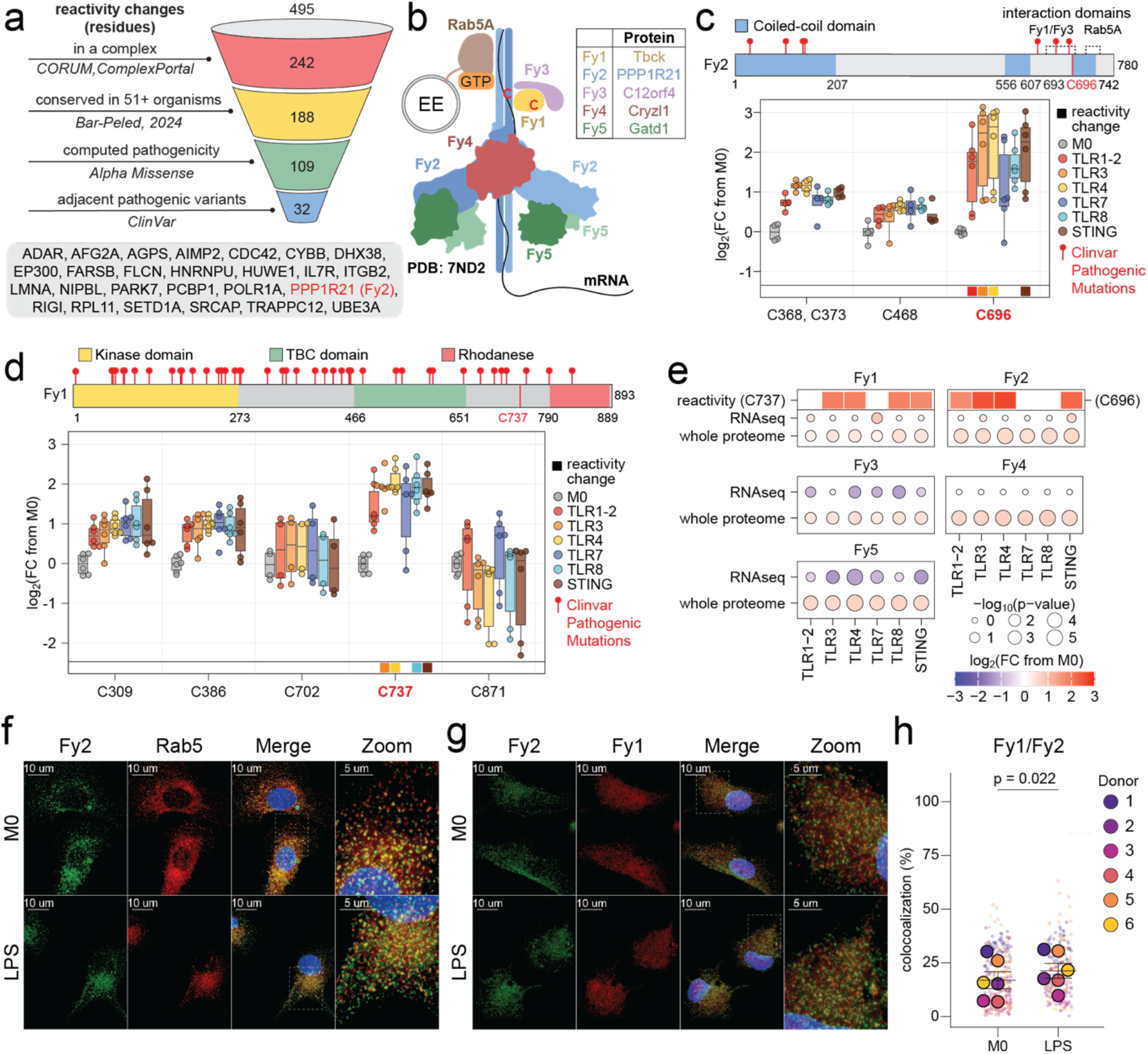
The RNA-trafficking FERRY complex assembles in macrophages and associates with early endosomes. **a**, Prioritization workflow for differentially reactive cysteines with potential functional relevance. Residues were sequentially filtered for significant reactivity changes in at least one macrophage stimulation condition (495 cysteines), localization within proteins annotated as members of protein complexes (CORUM^55, 56^ and Complex Portal^57, 58^; 242 cysteines), evolutionary conservation across 102 organisms^54^ (188 cysteines), predicted pathogenicity by AlphaMissense^53^ (109 cysteines), and proximity to pathogenic or likely pathogenic germline variants reported in ClinVar^59^ (within 15 residues; 32 cysteines). Top 32 cysteine-containing peptides mapped to 26 high-confidence protein targets, which are listed in the grey oval. **b**, Schematic of the five-membered FERRY complex^20^, with individual subunits shown in different colors. Differentially reactive cysteines in Fy1 and Fy2 are denoted as red “C” symbols. Inset lists the UniProt protein names corresponding to each FERRY complex member. **c**,**d**, Top, schematic of Fy1 and Fy2 proteins with functional domains and pathogenic ClinVar variants annotated. Bottom, box plots showing changes in Fy2 and Fy1 expression (triangles) and cysteine reactivity (dots) across macrophage stimulation conditions compared to M0 cells. Boxes indicate the 25th to 75th percentiles, and whiskers extend to 1.5x the interquartile range. Boxes on the bottom margin indicate cysteines classified as differentially reactive. **e**, Multi-omic characterization of stimulation-dependent changes in FERRY complex members. For each subunit (Fy1-Fy5), dots show the change across the profiled stimulation conditions in transcript abundance (RNA-seq) and protein abundance (TMT-exp). Cysteine reactivity changes for the indicated residues are shown as colored boxes for proteins with reactivity changes. Dot color indicates the direction and magnitude of change, and dot size represents -log_10_(p-value). **f**,**g**, Representative confocal microscopy images of Fy2 (red) and Rab5 (green; early endosome marker) (**f**) or Fy2 (red) and Fy1 (green) (**g**) in M0 and TLR4-stimulated macrophages. Gray boxes highlight zoomed-in regions. **h**, Quantification of colocalized intensity of Fy1 with Fy2. The average across cells for each donor is represented by a larger dot. Lines represent mean and SEM for n = 6 donors. Significance was determined by a two-tailed paired t-test on the donor means.

This analysis identified 32 differentially reactive cysteines in 26 high-confidence protein targets, including PPP1R21, also known as Fy2, a subunit of the recently described FERRY complex^20, 21^ (Fig. 4a-d, Extended Data Fig. 6c). The FERRY complex is a five-membered Rab5 effector originally characterized in neurons, where it associates with early endosomes to facilitate mRNA transport, subcellular localization, and localized translation^20, 21^ (Fig. 4b). More recently, Fy2 has been proposed to form a three-subunit complex with Fy4 and Fy5 that mediates mRNA transport more broadly across the endo-lysosomal system^60, 61^. Three of the five FERRY subunits detected in our TMT-exp dataset, Fy1, Fy2, and Fy4, showed coordinated post-transcriptional upregulation of protein expression following macrophage activation (Fig. 4e).

TMT-ABPP revealed stimulation-dependent reactivity changes of both Fy1_C737 and Fy2_C696 (Fig. 4c-e). Fy2 serves as the central scaffold of the FERRY complex through interactions with mRNA cargo, Fy1, Fy3, Fy4, Fy5, and GTP-bound Rab5^20^. Notably, the differentially reactive Fy2_C696 lies in the established Fy1/Fy3 interaction interface at the *C-* terminal domain (residues 646-705)^20^, suggesting that its increased reactivity may reflect stimulation-induced remodeling of the FERRY complex. While structural information is currently unavailable for Fy1, C737 lies on the same tryptic peptide as a reported phosphotyrosine pY732^62^, raising the possibility of post-translational regulation.

While the FERRY complex has primarily been studied in neurons, where it mediates mRNA transport required for axonal function, localized translation in macrophages has been reported for only a limited number of transcripts, including actin mRNA at phagocytic cups^63^. To determine whether the complex is assembled in macrophages, we performed IF analysis of Fy1, Fy2, Fy4, and Rab5 in resting and activated cells (Fig. 4f-h, Extended Data Fig. 6d,e; Data S4). All FERRY subunits displayed punctate localization consistent with membrane-associated compartments. Rab5 and Fy2 showed approximately 50% colocalization, consistent with the reported localization of Fy2 to early endosomes. Fy4 and Fy2 also showed approximately 50% colocalization (Extended Data Fig. 6d,e). In contrast, Fy1 showed only approximately 20% colocalization with Fy2 under basal conditions, consistent with the established functions of Fy1 in other signaling pathways^60^. The colocalization of Fy2 with Rab5 remained unchanged following TLR4 stimulation, suggesting that these proteins are constitutive interactors in our model. By comparison, colocalization of both Fy1 and Fy4 with Fy2 increased moderately following TLR4 stimulation (Fig. 4h and Extended Data Fig. 6e). These results support a model in which Fy2 is constitutively associated with Rab5, while recruitment of other complex members may be context-dependent. This interpretation is consistent with previous evidence for state-dependent recruitment of Fy1 and Fy3 to a stable three-membered complex between Fy2, Fy4, and Fy5^61^.

### Orthogonal profiling defines the FERRY interactome and identifies DISC1 as a macrophage Fy2-interacting protein

To validate state-dependent FERRY complex assembly and identify macrophage-specific interacting proteins, we performed complementary immunoprecipitation-mass spectrometry (IP-MS) and imaging analyses (Fig. 5a). IP-MS of Fy1 and Fy2 confirmed enrichment of core FERRY subunits in both M0 and TLR4-stimulated macrophages (Fig. 5b,c and Extended Data Fig. 7a-d; Data S2-7), with the exception of Fy5, which was detected only in Fy2 pulldowns (Extended Data Fig. 7b-c). Fy2 IP-MS showed increased enrichment of Fy1 and Fy3 following TLR4 stimulation (Fig. 5c), consistent with the IF colocalization data and the proposed state-dependent recruitment of these complex members. Reciprocally, Fy1 IP-MS showed increased enrichment of Fy2 and Fy4 in TLR4-stimulated macrophages, providing further support for this model (Extended Data Fig. 7d).

**Fig. 5.**
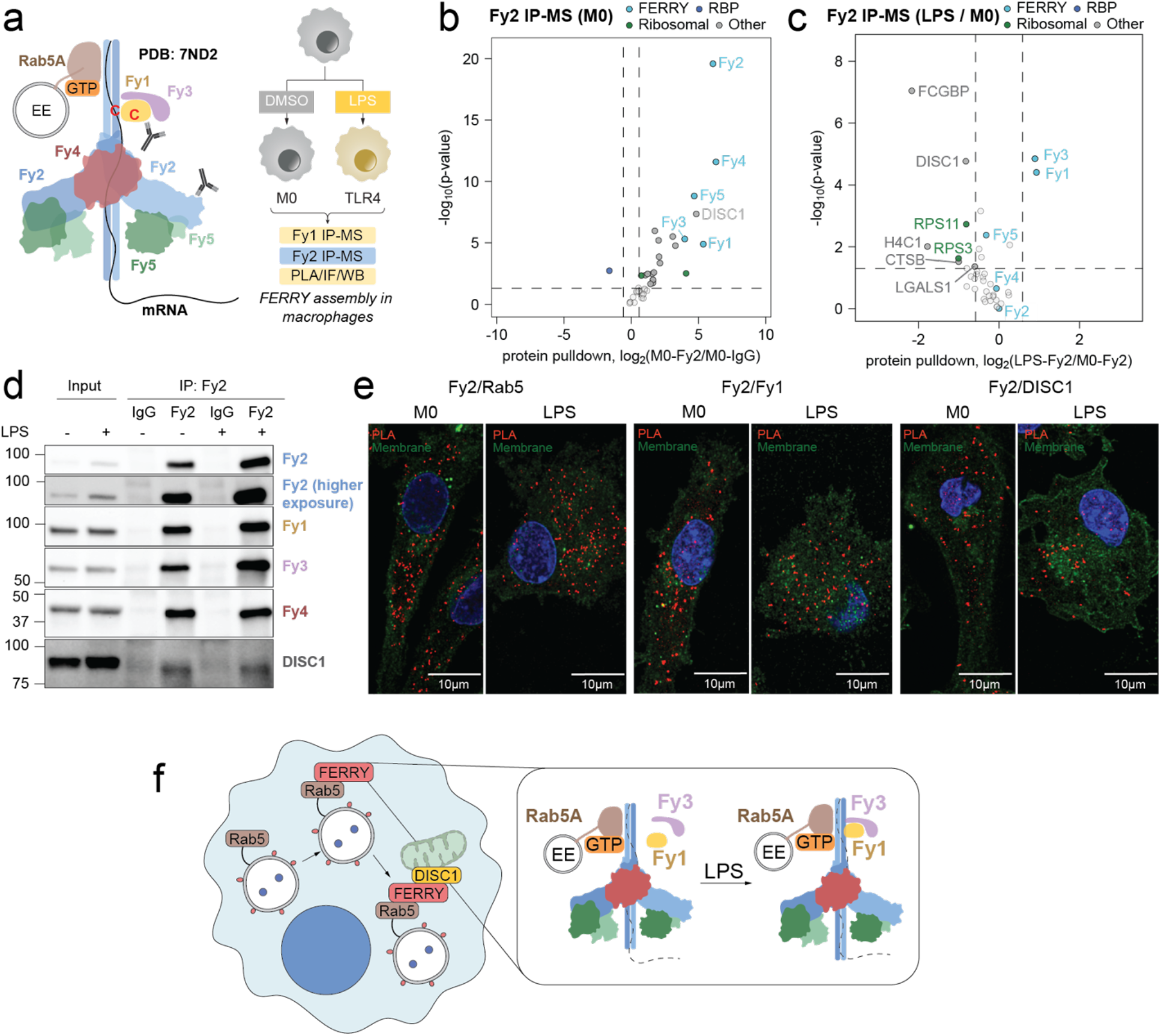
Characterization of state-dependency of FERRY complex assembly and composition. **a**, Schematic of experiments performed to investigate state-dependent differences in FERRY complex assembly and composition. Antibody symbols denote proteins targeted for IP-MS experiments. **b**, Volcano plot of CRAPome-filtered^64^ proteins co-enriched with endogenous Fy2 relative to an IgG control following immunoprecipitation from unstimulated M0 macrophages. Vertical dashed lines mark the fold-change cutoff (±log₂(1.5)) and the horizontal dashed line marks p = 0.05. Proteins exceeding both thresholds in the upper right quadrant are considered significantly enriched. Points are colored by protein category: FERRY complex subunits (light blue); RNA-binding proteins (RBPs; Gerstberger et al., 2014^65^; dark blue); ribosomal proteins (GO annotation; green); and all other proteins (grey). **c**, Volcano plot comparing CRAPome-filtered proteins co-enriched with endogenous Fy2 following immunoprecipitation from M0 and TLR4-stimulated macrophages. **d**, Representative Western blots showing Fy2 and co-immunoprecipitated Fy1, Fy3, Fy4, and DISC1 following immunoprecipitation of endogenous Fy2 from M0 and TLR4-stimulated macrophage lysates. **e**, Representative PLA images detecting interactions between Fy2 and Rab5 (left), Fy2 and Fy1 (middle), or Fy2 and DISC1 (right) in M0 and TLR4-stimulated macrophages. PLA signals are shown in red, and cell membranes and nuclei are shown in green and blue, respectively. **f**, Proposed model of state-dependent remodeling of the FERRY complex following TLR4 stimulation in macrophages.

The Fy1 interactome was substantially broader than that of Fy2, suggesting that Fy1 may have functions beyond its role in the FERRY complex, potentially mediated by its TBC domain which is characteristic of RabGAP proteins^66^. Consistent with this, gene ontology analysis of molecular functions enriched among Fy1-interacting proteins identified cytoskeletal binding, motor activity, and GTPase binding (Extended Data Fig. 7e). Considering the breadth of the Fy1 interactome and the central scaffolding role of Fy2 within the FERRY complex, we focused subsequent analyses on Fy2-associated proteins.

We next compared our combined macrophage Fy2 interactome with FERRY interactomes generated by GST-FERRY IP-MS from HEK293 cell lysates^21^ and Fy1/2 affinity purification mass spectrometry (AP-MS) in HeLa cells^60^. All five core FERRY subunits were consistently enriched across all datasets (Extended Data Fig. 7f,g). This comparison also identified candidate macrophage-specific interacting proteins, including the scaffolding protein DISC1, which associated with Fy2 in both resting and TLR4-stimulated macrophages but was not detected previously. We confirmed enrichment of core complex members and DISC1 by Western blot (Fig. 5d). Finally, to assess FERRY complex assembly in intact cells, we performed proximity-ligation assays (PLA), which report physical proximity of two proteins as discrete fluorescent puncta (Fig. 5e and Extended Data Fig. 7h,i). Analysis of PLA results supported the proximity of Fy2 to Rab5, Fy1, Fy4, and DISC1 in primary human macrophages, further validating FERRY complex assembly and its association with DISC1.

Combined, our findings support a model in which FERRY assembles and associates with early endosomes in primary human macrophages, with TLR4 stimulation promoting post-translational recruitment of Fy1 and Fy3 to the complex (Fig. 5f). We further identify DISC1 as a previously unrecognized Fy2-interacting protein. Future studies will define the role of DISC1 in FERRY complex function and determine how remodeling of the complex contributes to macrophage activation.

## DISCUSSION

Macrophages are central effectors of innate immunity and contribute to the pathogenesis of inflammatory and autoimmune diseases and cancer. Activation of cell-surface and endosomal TLRs, as well as cytosolic DNA-sensing cGAS-STING pathway, induces inflammatory programs that have been extensively characterized at the transcriptional level. However, macrophage activation also involves rapid post-translational regulation that governs protein abundance, activity, localization, and molecular interactions to remodel metabolism, organelle function, and membrane trafficking. The scope of this regulatory layer remains incompletely defined, in part because conventional proteomic approaches do not capture the full spectrum of biochemical changes associated with macrophage activation that occur independently of RNA or protein abundance changes.

Here, we integrated transcriptomics, global proteomics, phosphoproteomics, and PTM-agnostic, multiplexed cysteine reactivity profiling by TMT-ABPP to map the post-translational landscape of primary human macrophages following TLR and STING activation. These complementary approaches revealed substantial discordance between transcript- and protein-abundance changes, as well as extensive post-translational remodeling not captured by abundance measurements alone. Proteins exhibiting abundance changes in the absence of corresponding transcriptional regulation were enriched in metabolic pathways, mitochondrial gene expression, and mitochondrial translation – processes that support macrophage activation and require rapid remodeling in response to extracellular cues. Our analyses further identified differential protein turnover as a potential contributor to the observed divergence between transcript- and protein-level changes, although observed discordance could also be due to other factors such as protein translation rates and stabilizing interactions. These findings highlight the contributions of post-transcriptional mechanisms to proteome remodeling during macrophage activation.

Cysteine reactivity profiling provided an orthogonal readout of changes in native protein states. Cysteine labeling by electrophilic probes can be influenced by local structure, ligand occupancy, protein interactions, subcellular environment, and reversible or irreversible covalent modifications. Accordingly, differential reactivity does not by itself identify a specific regulatory mechanism, but it can nominate protein sites undergoing biochemical remodeling that would not be detected by conventional unenriched proteomics or PTM-focused analyses. Reactivity changes observed following macrophage activation showed limited overlap with those identified in our previous studies of T cell activation^15^ and dysfunction^18^. Although differences in protein coverage and stimulation conditions may contribute to this limited overlap, the results suggest that cysteine reactivity profiling can capture distinct biochemical states across immune cell populations.

Cysteine reactivity profiling revealed extensive remodeling of small-GTPase signaling and vesicular trafficking pathways, including endosomal transport and autophagy. These processes are central to macrophage biology, controlling receptor trafficking, phagocytosis, cytokine secretion, antigen processing, and organelle homeostasis. Among these changes, we identified altered cysteine reactivity in the nucleotide-binding pocket of CDC42, an important GTPase with context-dependent functions in macrophages^67, 68, 69, 70^ regulating actin remodeling, cell migration, phagocytosis, and inflammatory signaling. TLR4 stimulation has been reported to activate CDC42 in macrophage models^71^. In lysate-based metabolite-addback experiments, GTP, but not ATP, decreased CDC42_C18 labeling by a cysteine-reactive maleimide probe, with a more pronounced effect in basal state M0 macrophages compared to TLR4-stimulated cells. Whole-cell GTP abundance did not change significantly following macrophage activation, indicating that differential CDC42_C18 reactivity is unlikely to be explained by changes in bulk intracellular GTP concentration, but may instead reflect targeted regulation of CDC42 by GEFs and GAPs that affect nucleotide occupancy, protein conformation, or effector protein interactions. It is also worth noting that CDC42_C18 is located within a redox-sensitive motif conserved among several Rho-family GTPases, raising the possibility that reactivity could also be influenced by oxidative modifications, including *S-*nitrosylation or sulfenylation^72, 73, 74^. While the current data do not distinguish among these mechanisms, it nevertheless illustrates how cysteine reactivity profiling can detect regulated changes at functional protein sites that are not apparent from metabolite or protein abundance measurements alone.

Integrated analysis of structural, evolutionary, and disease-associated annotations further prioritized cysteines in the FERRY complex as candidate sites of biochemical or post-translational regulation in primary human macrophages. FERRY is a Rab5 effector that associates with early endosomes and select mRNAs, enabling endosome-dependent mRNA transport and localized translation. We identified activation-associated reactivity changes in two subunits of this five-membered complex: the central scaffold Fy2 (PPP1R21) and its direct interactor Fy1 (TBCK)^21^. While the FERRY complex has been characterized in neurons and other cell lines, its assembly and function in immune cells has remained largely unknown. We validated complex assembly and localization to early endosomes in primary human macrophages using complementary biochemical, proteomic, and imaging approaches, including IP-MS of both Fy1 and Fy2, immunofluorescence, and proximity ligation assays. These findings extend FERRY beyond the systems in which it was originally characterized and raise the possibility that endosome-associated mRNA transport contributes to innate immune cell function.

Our IP-MS analysis further identified DISC1 as a previously unrecognized Fy2 interactor, which we validated by Western blot analysis following Fy2 immunoprecipitation, and by PLA. DISC1 is a multifunctional scaffolding protein that coordinates microtubule-dependent transport, including mRNA and mitochondria trafficking^75^, but has not been previously linked to FERRY. FERRY preferentially associates with subsets of transcripts, including mRNAs encoding mitochondrial proteins, and mRNA-loaded FERRY-positive endosomes have been observed in close proximity to mitochondria^20, 21^. Recent work has also supported a potential role of the dynein-dynactin adaptor JIP4 in the endo-lysosomal FERRY trafficking^61^. In this context, the identified Fy2-DISC1 association suggests that DISC1 may help coordinate the movement or positioning of FERRY-positive endosomes relative to mitochondria, potentially promoting spatially restricted delivery or translation of mitochondrial transcripts. Further studies will be required to define the molecular basis and functional significance of this interaction during macrophage activation.

Localized translation remains poorly understood in immune cells. During innate immune activation, macrophages remodel their endosomal and lysosomal compartments, cytoskeleton, secretory pathways, and metabolic state. Coupling select transcripts to motile endosomes could allow protein synthesis to be coordinated with these spatially restricted changes. Our findings raise the possibility that FERRY participates in this process, but its RNA cargoes and functional outputs in macrophages remain unknown. Identifying FERRY-associated transcripts in resting and activated macrophages, determining whether they are translated near endosomes or other organelles, and perturbing individual FERRY subunits will be necessary to establish whether this pathway regulates inflammatory signaling, organelle homeostasis, or other macrophage functions.

Several limitations should be considered. Our study focused on a 24-hour endpoint and therefore doesn’t capture the dynamic regulatory events that occur during earlier stages of macrophage activation. Moreover, cysteine reactivity changes can arise through multiple mechanisms and require site-specific biochemical validation before they can be assigned to changes in protein activity or a particular post-translational modification. Similarly, metabolite addback experiments performed in lysates do not preserve subcellular compartmentalization of metabolite pools and regulatory layers present in intact cells. Finally, our FERRY studies establish complex assembly, localization, and its association with DISC1 in macrophages but do not yet define its RNA cargoes or functional contribution to macrophage activation. Addressing these questions will require additional investigations that extend beyond the current study.

In summary, our integrated multi-omic analysis provides a comprehensive view of biochemical and post-translational changes accompanying TLR and STING activation in primary human macrophages. The data identify extensive remodeling of small-GTPase signaling and the endo-lysosomal system, nominate functional cysteines that may report on changes in nucleotide occupancy and other regulatory states, and establish the presence and assembly of the FERRY mRNA transport complex in macrophages. These datasets provide a rich resource for identifying regulatory nodes of macrophage activation and lay the groundwork for future investigations into the role of mRNA transport and localized translation in innate immunity.

## Acknowledgements

We thank Nacim Kerrouche for preliminary investigations. T.L.Z., H.S., H.K., and E.V.V. were supported by The Stavros Niarchos Foundation Institute for Global Infectious Disease Research at Rockefeller and CZ Biohub New York Investigator Program. T.L.Z. was also supported by NIH (T32GM136640-Tan to T.L.Z). H.S., H.K., and E.V.V. were supported by The Alexandrine and Alexander L. Sinsheimer Fund and Herbert and Nell Singer Foundation. B.Z. was supported by NIH 1F31CA317812. J.B. was supported by a career development award from Blood Cancer United, the Lauri Strauss Leukemia Research Foundation and the MPN Research Foundation. O.A.-W. is supported by the Neil S. Hirsch Foundation, Edward P. Evans Foundation, Break Through Cancer, NIH/NCI (R01 CA251138, R01 CA242020, R01 CA283364, and P50 CA254838), Cancer Center Support Grant (P30 CA008748), NIH/NHLBI (R01 HL128239), and the Leukemia & Lymphoma Society. E.V.V. was also supported by Rockefeller University start-up funds, Robertson Foundation, The Achelis and Bodman Foundation, Weill Cancer Hub East, and Searle Scholar Program. H.K. and A.v.V. were supported by Pels Family Center Fellowships. A.v.V. was also supported by a fellowship from National Cancer Center. C.R.W. was supported by the Marlene Hess Center for Research on Women’s Health and Biomedicine and Kavli NSI Postdoctoral Fellowship. Confocal microscopy, image processing and analysis were performed in The Rockefeller University’s Bio-Imaging Resource Center (RRID:SCR_01779). Flow cytometry assays were optimized with guidance from Dr. Svetlana Mazel and the Rockefeller Flow Cytometry Resource Center (RRID:SCR_017694). Some analyses were performed on computational resources from the Rockefeller University High Performance Computing Resource Center (RRID: SCR_025889).

## Author contributions

Conceptualization: T.L.Z., E.V.V.

Data curation: T.L.Z., H.S., B.Z., E.V.V.

Formal analysis: T.L.Z., H.S., Y.K., P.B., E.V.V.

Investigation: T.L.Z., R.G., A.v.V., H.K., C.R.W., J.B., E.V.V.

Methodology: T.L.Z., R.G., A.v.V., H.K., C.R.W., E.V.V.

Validation: T.L.Z., R.G., A.v.V., H.K., C.R.W., H.S., E.V.V.

Resources: M.Z., E.V.V.

Software: H.S.

Visualization: T.L.Z., A.v.V., C.R.W., Y.K., H.S., E.V.V.

Supervision: O.A.-W., M.Z., E.V.V.

Writing – original draft: T.L.Z., H.S., E.V.V.

Writing – review & editing: all authors read, edited, and approved the final manuscript

## Competing interests

E.V.V. is listed as a co-inventor on patents with Vividion Therapeutics. J.B. has previously served on an advisory board for Sobi (Swedish Orphan Biovitrum). O.A.-W. is a founder and scientific advisor of Codify Therapeutics, holds equity and receives research funding from this company. O.A.-W. has served as a consultant for Amphista Therapeutics, and MagnetBio, and is on scientific advisory boards of Envisagenics Inc. and Harmonic Discovery Inc.; O.A.-W. received research funding from Astra Zeneca, Nurix Therapeutics, and Minovia Therapeutics, unrelated to this study. The remaining authors declare no competing interests.

## Data, code, and materials availability

- Raw proteomic data and RNA-sequencing data will be deposited to PRIDE and NCBI GEO.
- The code used for data processing and visualization will be deposited to GitHub.
- Any additional information required to reanalyze the data reported in this paper is available from the lead contact upon request.

## Extended Data Figures

**Extended Data Fig. 1.**
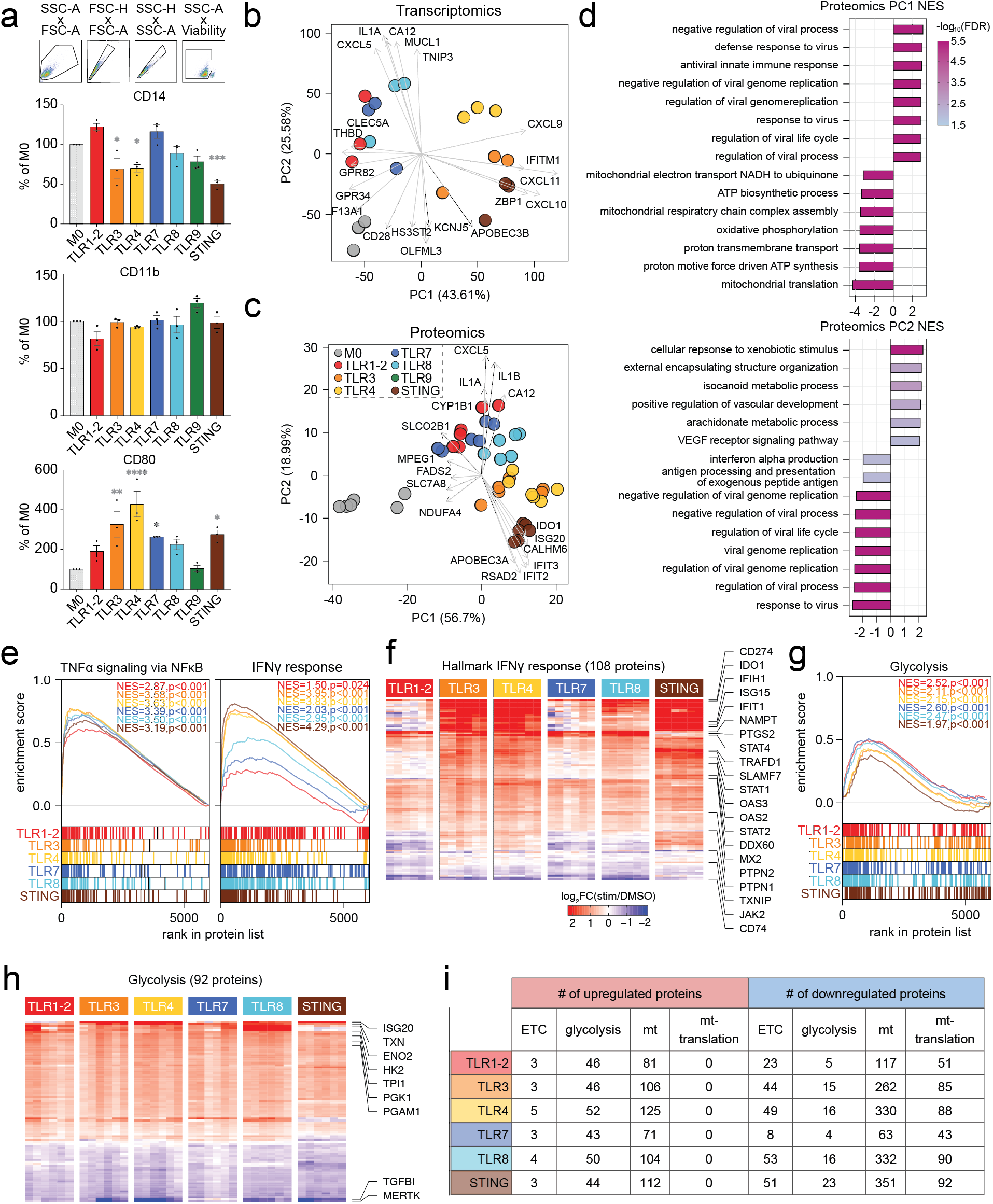
Multi-omic profiling of TLR- and STING-stimulated macrophages reveals induction of pro-inflammatory pathways (Related to Figure 1). (**a**) Representative flow cytometry gating strategy (top) and quantification of cell-surface expression of CD14, CD11b, and CD80 across macrophage activation states (bottom). Data are presented as the average ± SEM of three donors. Statistical significance was determined by one-way matched ANOVA with Dunnett’s multiple comparisons test (ns, p > 0.05; *, p < 0.05; **, p < 0.01; ***, p < 0.001; ****, p < 0.0001). (**b**) Principal component analysis (PCA) of bulk transcriptomic data from n = 3 donors. Normalized count values were log_2_ transformed, and the 5,000 protein-coding genes with the highest cross-experiment variability were used. The five highest and five lowest loadings from PC1 and PC2 were plotted. (**c**) PCA of unenriched proteomic data from n = 3 donors. Protein abundance ratios were log_2_ transformed and only proteins quantified in all experiments were used. The five highest and five lowest loadings from PC1 and PC2 were plotted. (**d**) Gene set enrichment analysis (GSEA) of PCA loading scores using the Gene Ontology Biological Process database. Top 15 gene sets based on normalized enrichment scores (NES) are plotted for PC1 (top) and PC2 (bottom). (**e**) GSEA of the Hallmark TNFα signaling via NFκB and IFNγ response gene sets. Each stimulation condition was compared to unstimulated M0 macrophages. NES, normalized enrichment score. (**f**) Heatmap displaying abundance changes for proteins in the Hallmark IFNγ response gene set relative to M0 cells. Proteins were hierarchically clustered. (**g**) GSEA of the Hallmark Glycolysis list. Each stimulation condition was compared to M0 cells. NES, normalized enrichment score. (**h**) Heatmap displaying abundance changes for proteins in the Hallmark Glycolysis gene set relative to M0 cells. Proteins were hierarchically clustered. (**i**) Numbers of upregulated and downregulated proteins in each macrophage activation state associated with GO terms ETC, Glycolysis, Mitochondria, and Mitochondrial translation.

**Extended Data Fig. 2.**
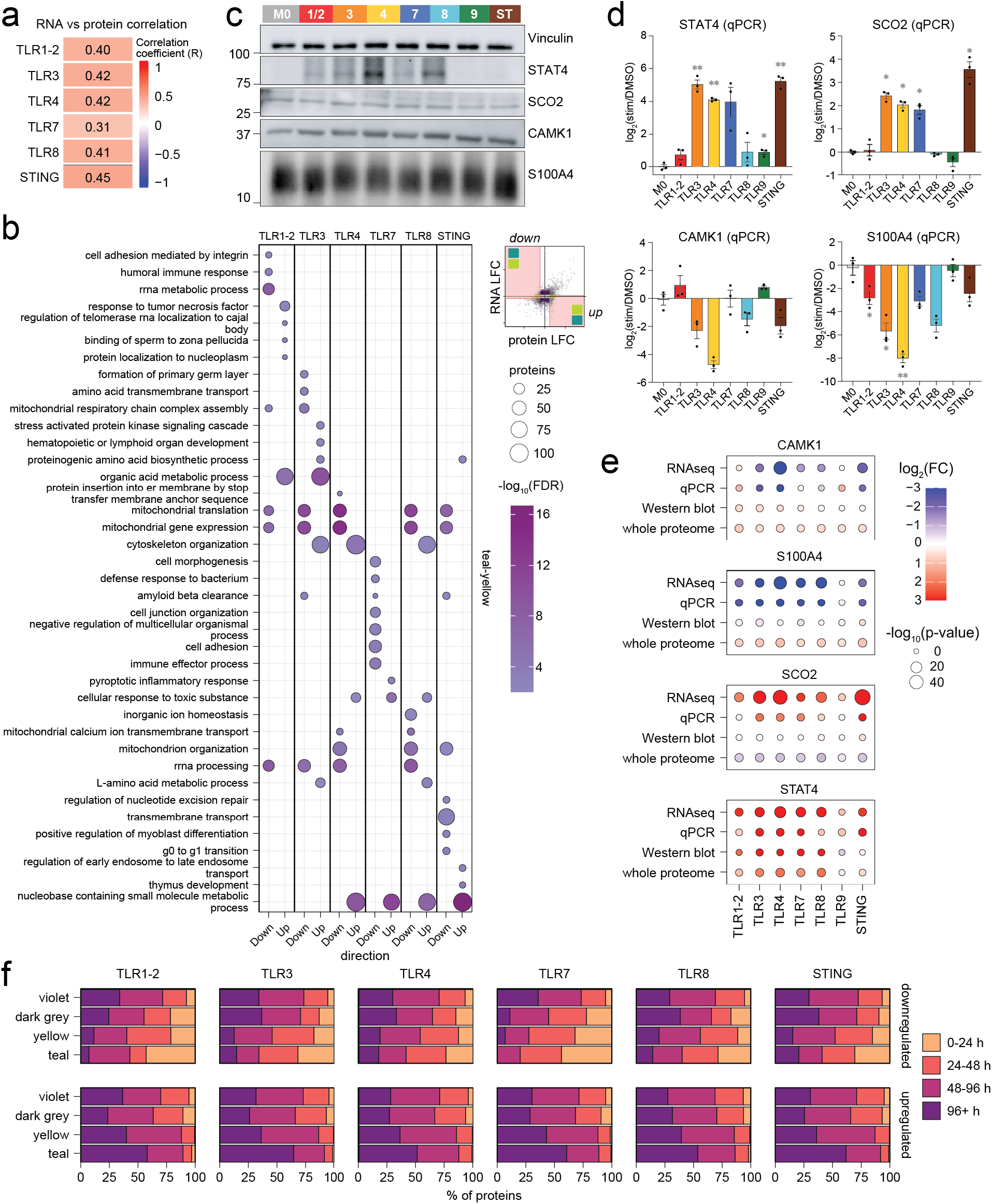
Characterization of genes with discordant protein- and transcript-level changes (Related to Figure 1). (**a**) Heatmap of Pearson’s Correlation Coefficients comparing protein- and transcript-level log_2_ fold changes (LFC) across macrophage stimulation conditions compared to unstimulated M0 cells. (**b**) Overrepresentation analysis (Fisher’s exact test, experiment-specific background) of proteins changing in abundance without corresponding transcriptional changes across macrophage stimulation conditions compared to M0 cells (yellow and teal quadrants in Fig. 1h). Terms with FDR < 0.01 and protein number > 5 are shown. Reg., regulation; Up, positive protein LFC; Down, negative protein LFC. (**c**) Representative Western blots of selected proteins showing concordant or discordant protein- and transcript-level changes following macrophage stimulation. (**d**) RT-qPCR quantification of transcript levels for selected genes. Data shown represents average ± SEM from three donors, and statistical comparison is by one-way matched ANOVA with Dunnett’s multiple comparisons test (ns, p > 0.05; *, p < 0.05; **, p < 0.01). (**e**) Summary of transcript-level (RNA-seq and RT-qPCR) and protein-level (Western blot and abundance-based proteomics) changes in STAT4, SCO2, CAMK1, and S100A4 across macrophage activation conditions compared to M0 cells. p-values were calculated depending on data modality: DESeq2 for RNA-Seq, ANOVA for RT-qPCR, one sample t-test (against 1, the value M0 is normalized to) for Western blotting, and unpaired t-test for TMT-exp (consistent throughout the manuscript). Fold-change values represent the mean for each modality. (**f**) Reference protein half-lives in monocytes^1^ for upregulated and downregulated proteins, depending on their correlation with transcript-level changes. Half-life values represent the mean of two biological replicates, and only proteins with reported half-life data are included.

**Extended Data Fig. 3.**
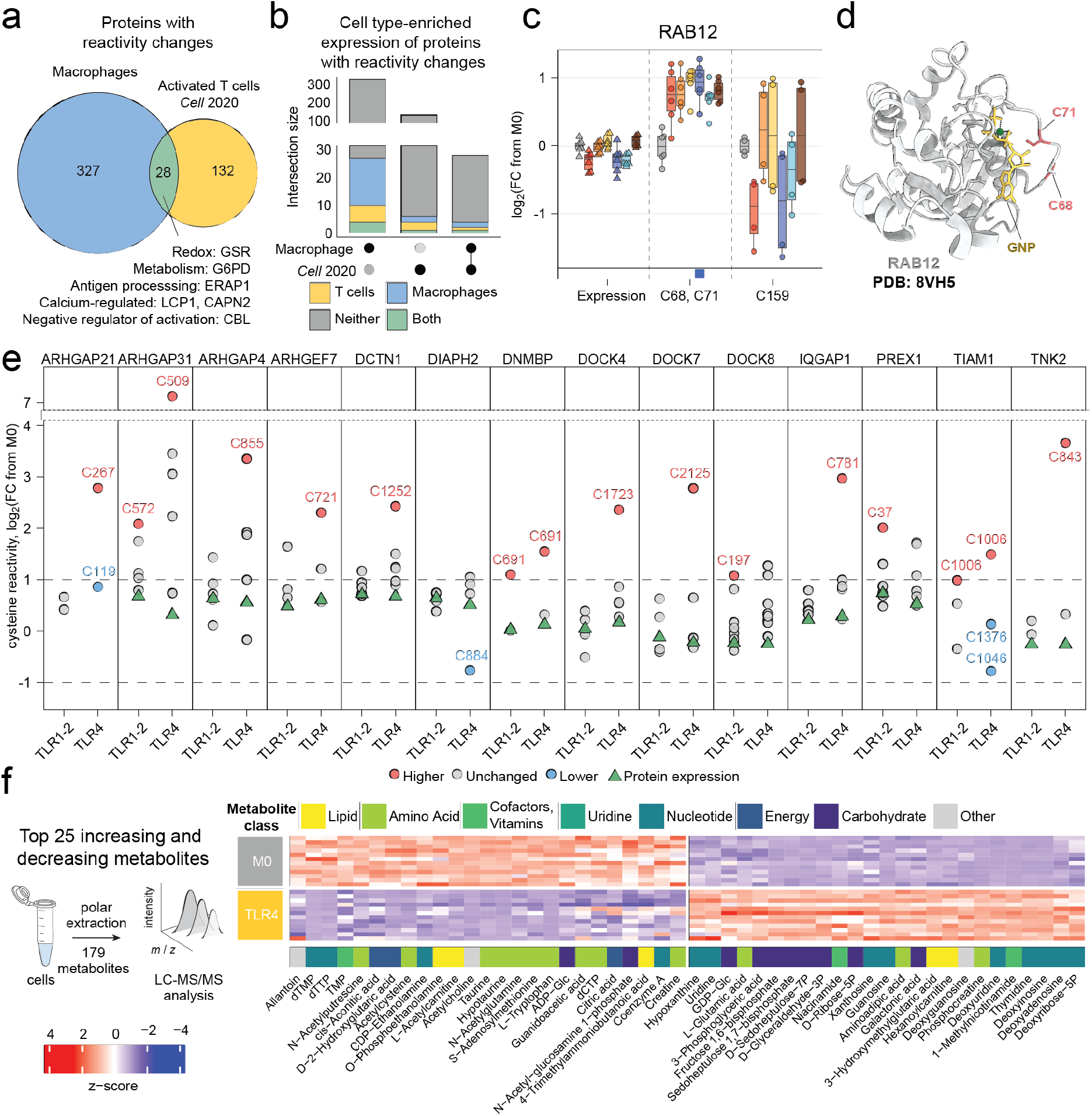
Cysteine reactivity profiling and metabolomics reveal remodeling of GTPases and their regulatory networks (Related to Figure 2). (**a**) Venn diagram comparing proteins with reactivity changes identified in TLR- and STING-stimulated primary human macrophages (this study; blue) with those identified in TCR-stimulated primary T cells^2^. Select proteins in the intersection are highlighted. (**b**) UpSet plot comparing proteins with cysteine reactivity changes in primary human macrophages and T cells^2^. Bars are colored according to cell type-enriched expression annotations from the Human Protein Atlas^3^, combining the "Cell type enhanced," "Cell type enriched," and "Group enriched" categories. (**c**) Box plots showing changes in RAB12 expression (triangles) and cysteine reactivity (dots) across macrophage stimulation conditions compared to M0 cells. Boxes indicate the 25th to 75th percentiles, and whiskers extend to 1.5x the interquartile range. Boxes on the bottom margin indicate cysteines classified as differentially reactive. (**d**) Cryo-EM structure of RAB12 bound to a GTP analog (PDB 8VH5^4^). Cysteines contained within the same differentially reactive tryptic peptide are highlighted in red. (**e**) Log_2_ fold changes in protein expression and cysteine reactivity for CDC42-interacting proteins in TLR1/2- and TLR4-stimulated macrophages relative to M0 cells. Protein expression changes are shown as green triangles and cysteine reactivity changes as dots. Cysteine reactivity and protein expression data are from n = 3 donors. (**f**) Heatmap showing z-scores calculated from metabolite signal intensity in M0 and TLR4-stimulated macrophages. Top 25 increasing and decreasing metabolites by p-value between TLR4-stimulated macrophages (bottom) and M0 cells (top) are shown. Annotations at the bottom indicate metabolite class. Data are from n = 4 donors.

**Extended Data Fig. 4.**
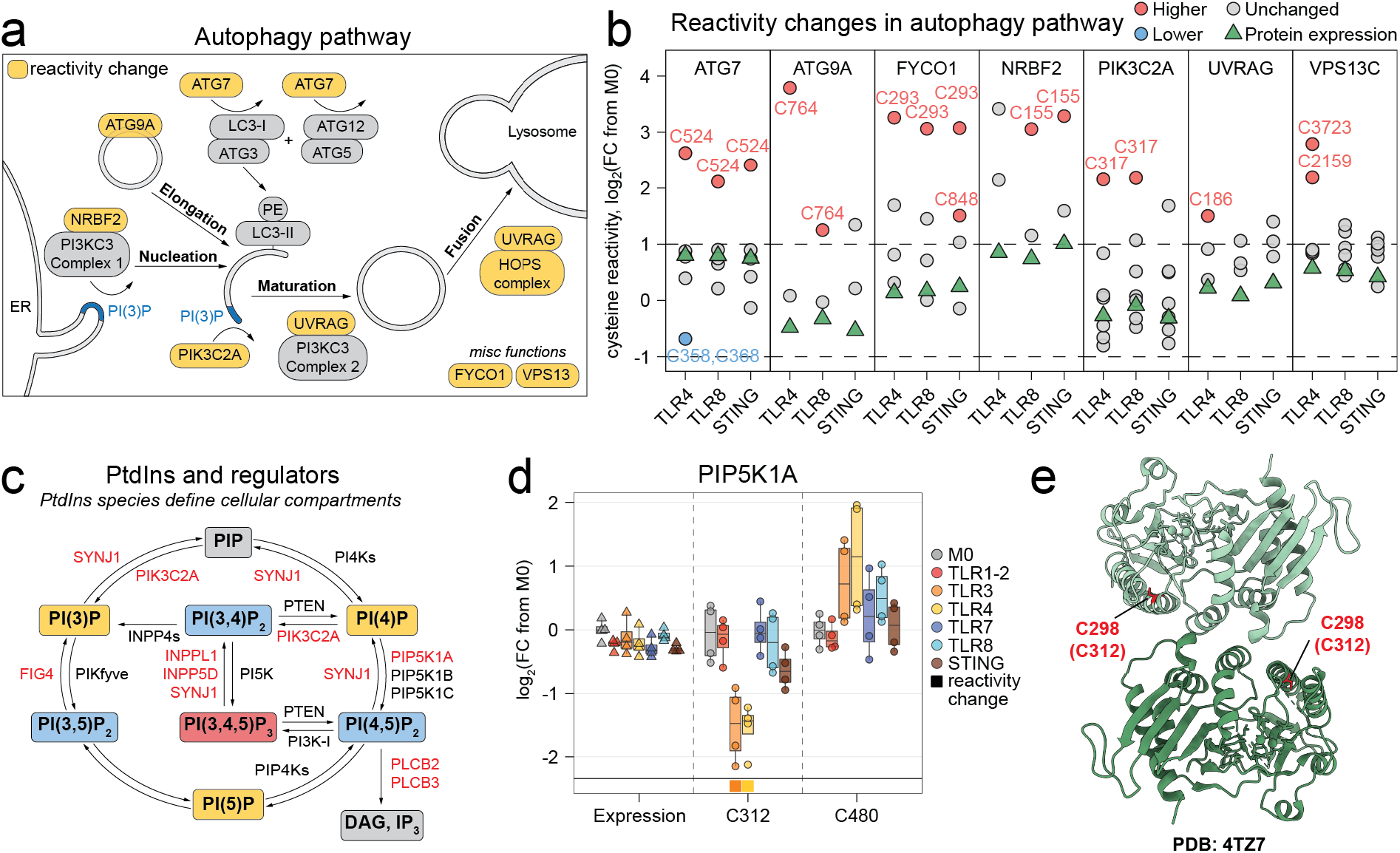
Cysteine reactivity profiling reveals remodeling of autophagy pathways and phosphoinositide regulators (Related to Figure 3). (**a**) Schematic of the autophagy pathway. Proteins with reactivity changes in any condition are highlighted in yellow. (**b**) Log_2_ fold changes in protein expression and cysteine reactivity for proteins in the autophagy pathway in TLR4-, TLR8-, and STING-stimulated macrophages relative to M0 cells. Protein expression changes are shown as green triangles and cysteine reactivity changes as dots. Cysteine reactivity and protein expression data are from n = 3 donors. (**c**) Schematic of phosphoinositide species and the lipid kinases and phosphatases that catalyze their interconversion. PtdIns containing one, two or three phospho groups are shown in yellow, blue, and red, respectively. Proteins with cysteine reactivity changes are highlighted in red. (**d**) Box plots showing changes in PIP5K1A expression (triangles) and cysteine reactivity (dots) across macrophage stimulation conditions compared to M0 cells. Boxes indicate the 25th to 75th percentiles, and whiskers extend to 1.5x the interquartile range. Boxes on the bottom margin indicate cysteines classified as differentially reactive. (**e**) Crystal structure of the zebrafish PIP5K1A dimer (PDB 4TZ7^5^), with the cysteine corresponding to the differentially reactive C312 in human PIP5K1A (C298 in zebrafish) highlighted.

**Extended Data Fig. 5.**
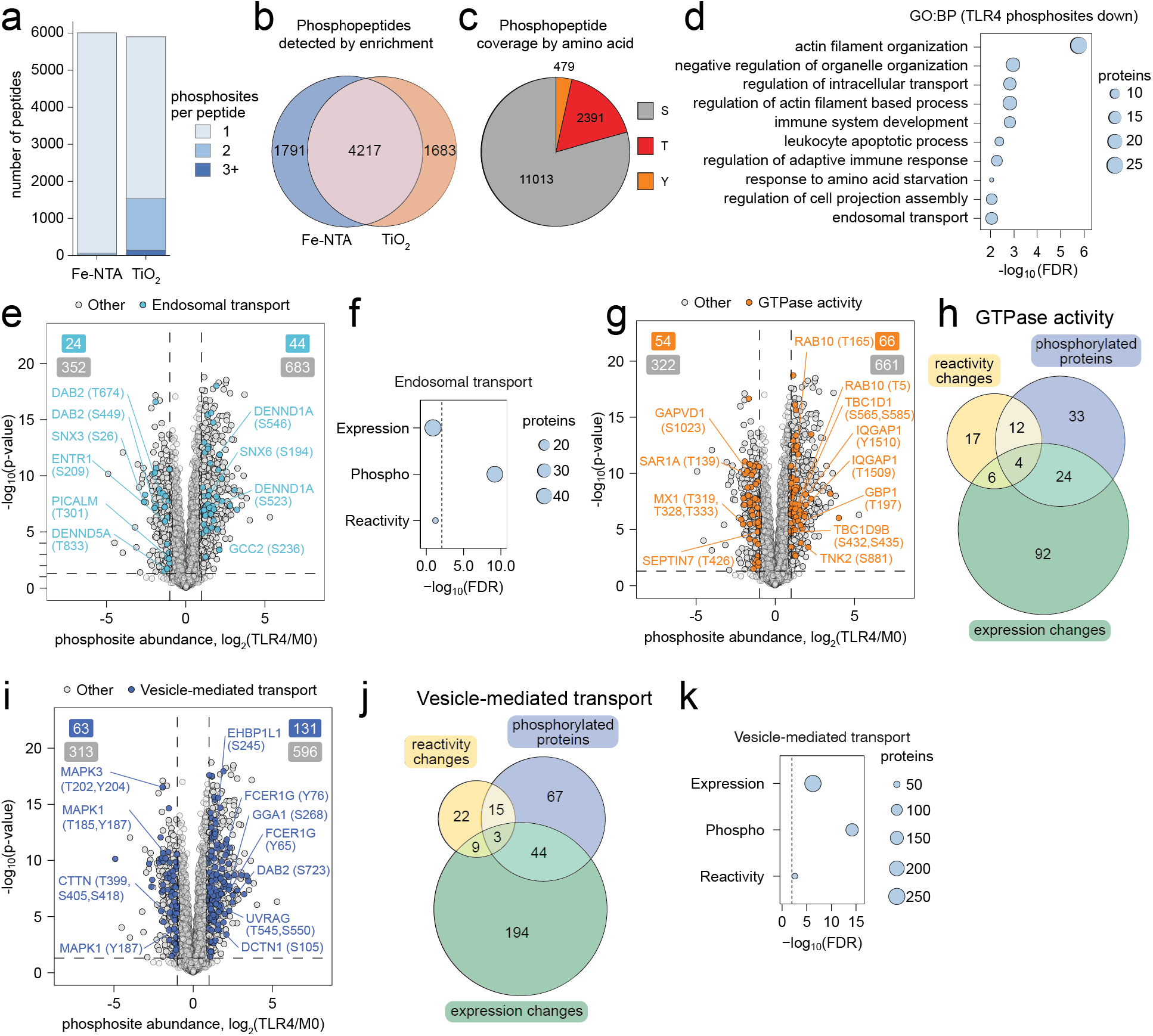
Phosphoproteomic profiling reveals enrichment of vesicle trafficking pathways in TLR4-stimulated macrophages (Related to Figure 3). (**a**) Phosphopeptides detected following Fe-NTA and TiO_2_ enrichment, colored by the number of phosphosites per peptide. Data are from n = 4 donors. (**b**) Venn diagram comparing phosphopeptides detected following Fe-NTA and TiO_2_ enrichment. (**c**) Pie chart showing the distribution of serine, threonine, and tyrosine phosphosites detected using the dual TiO_2_/Fe-NTA enrichment workflow. (**d**) GO:BP enrichment analysis of proteins containing phosphosites with significantly decreased phosphorylation in TLR4-stimulated cells compared to M0 (FC < 0.5, p < 0.05). Terms with FDR < 0.01 and protein number > 5 are shown. **(e)** Volcano plot showing log_2_ fold changes of phosphopeptides between TLR4-stimulated and unstimulated M0 cells where each point represents a phosphosite, with proteins annotated in the endosomal transport gene ontology term (GO:0016197) highlighted in teal. **(f)** Comparative enrichment of the endosomal transport gene set across expression proteomics, reactivity profiling, and phosphoproteomic datasets. **(g)** Volcano plot showing log_2_ fold changes of phosphopeptides between TLR4-stimulated and unstimulated M0 cells where each point represents a phosphosite, with proteins annotated in the GTPase activity gene ontology term (union of molecular function terms related to GTPase activity, activation, regulation, inhibition, and binding) highlighted in orange. **(h)** Venn diagram of the differentially regulated proteins identified by expression proteomics, cysteine reactivity profiling, and phosphoproteomic datasets within the GTPase activity gene set. **(i)** Volcano plot showing log_2_ fold changes of phosphopeptides between TLR4-stimulated and unstimulated M0 cells where each point represents a phosphosite, with proteins annotated in the vesicle-mediated transport gene ontology term (GO:0016192) highlighted in blue. **(j)** Venn diagram of the differentially regulated proteins identified by expression proteomics, cysteine reactivity profiling, and phosphoproteomic datasets within the vesicle-mediated transport gene set. (**k**) Comparative enrichment of the vesicle-mediated transport gene set across expression proteomics, reactivity profiling, and phosphoproteomic datasets. Gene sets for e-k were tested independently against the TLR4-regulated proteins identified by cysteine reactivity profiling (FC > 2 relative to expression value), expression-based proteomics (upregulated, FC > 1.5, p < 0.05), or phosphoproteomics (upregulated, FC > 2, p < 0.05) in TLR4-stimulated macrophages compared to M0 cells. Enrichment was computed with Fisher’s exact test against all annotated genes as background, and p-values were adjusted using the Benjamini-Hochberg method. The dashed line marks FDR = 0.01. Venn diagrams comparing proteins show changes in cysteine reactivity (FC > 2 relative to expression value), expression (FC > 1.5, p < 0.05), or phosphorylation (FC > 2, p < 0.05) in TLR4-stimulated macrophages compared to M0 cells.

**Extended Data Fig. 6.**
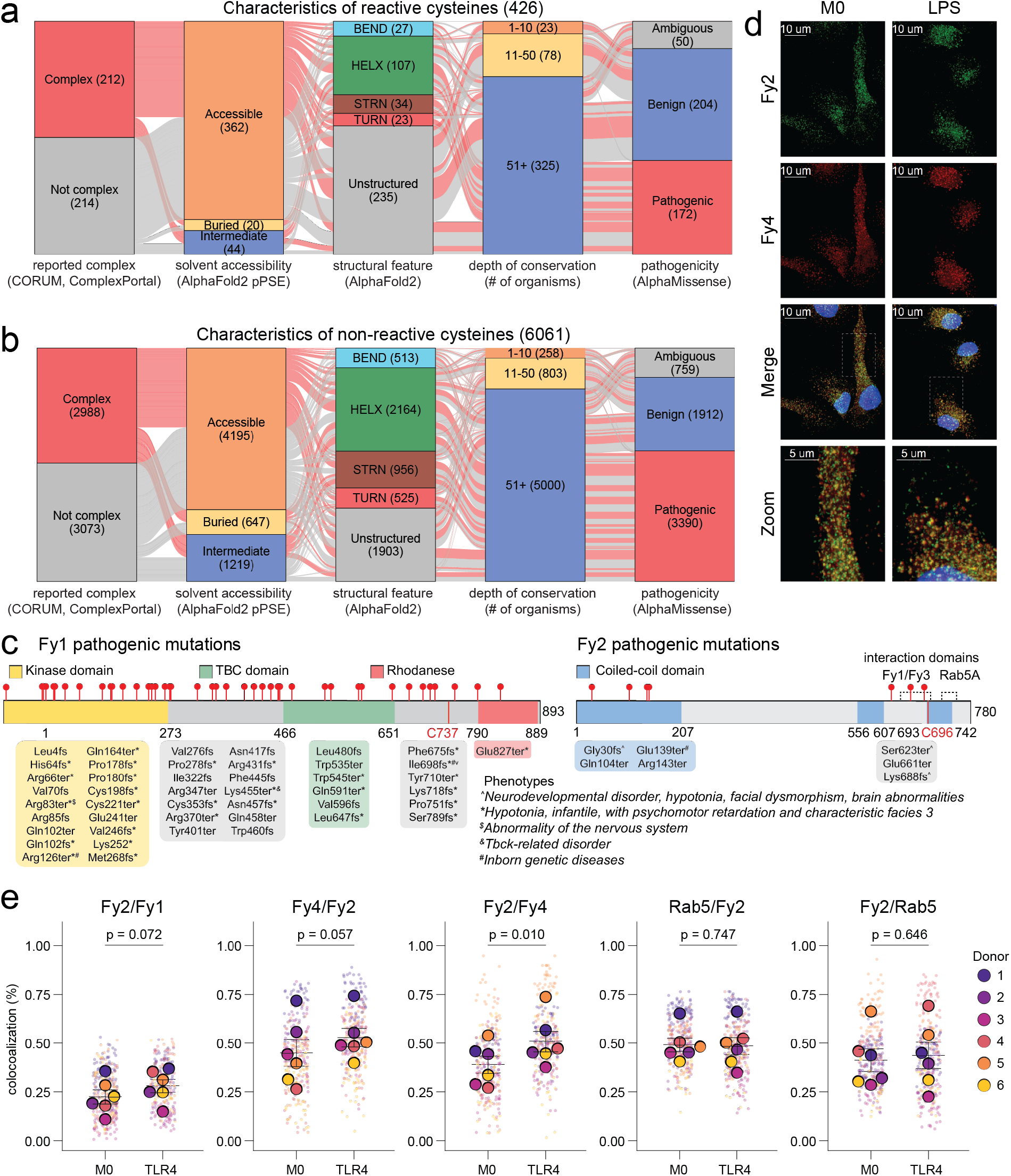
Prioritization of differentially reactive cysteines highlights assembly of the FERRY complex (Related to Figure 4). (**a, b**) Structural, evolutionary, and pathogenic features of differentially reactive cysteines compared with cysteines without reactivity changes. Alluvial diagrams partitioning (**a**) cysteines with significant reactivity changes (n = 399) and (**b**) cysteines without reactivity changes (n = 5,626) across five annotation categories: membership of the parent protein in a reported protein complex (CORUM) and Complex Portal; complex or not complex); predicted solvent accessibility based on AlphaFold2 structures (Buried = 9+ pPSE, Intermediate = 5-9 pPSE, Accessible = < 5 pPSE) local secondary structure (AlphaFold2/DSSP; helix [HELX], strand [STRN], turn [TURN], bend [BEND], or unstructured); evolutionary conservation, binned by the number of organisms in which the cysteine is conserved (1-10, 11-50, ≥51); and predicted variant pathogenicity (AlphaMissense classification: benign, ambiguous, or pathogenic). Each ribbon represents one cysteine. (**c**) ClinVar pathogenic or likely pathogenic mutations with a review status of at least one star mapped for Fy1 and Fy2. Variants are colored according to the protein domain they are in, with individual red markers denoting distinct variants. Unique markers denote corresponding phenotypes of each mutation. (**d**) Representative confocal microscopy images of Fy2 (red) and Fy4 (green) in M0 and TLR4-stimulated macrophages. White boxes highlight zoomed in regions. (**e**) Quantification of colocalized intensity between Fy1 and Fy2, Fy4 and Fy2, or Rab5 and Fy2. Average of cells per donor is represented by larger dot. Lines represent mean and SEM for n = 6 donors. Significance was determined by a two-tailed paired t-test on the donor means.

**Extended Data Fig. 7.**
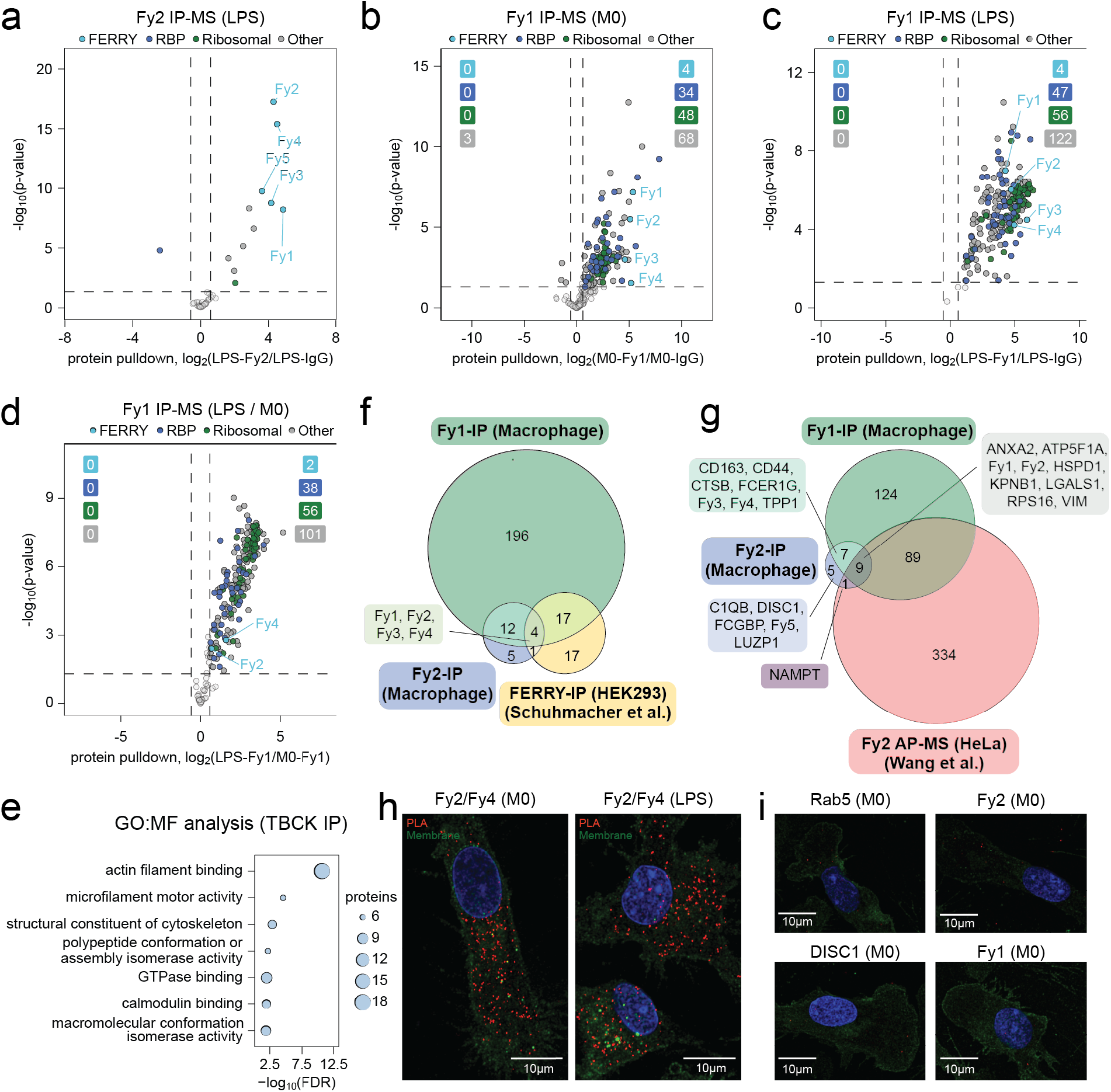
IP-MS and PLA reveal constitutive assembly of the FERRY complex, state-dependent recruitment of Fy1 and Fy3, and additional Fy1- and Fy2-interacting partners (Related to Figure 5). (**a-d**) Volcano plots of CRAPome-filtered^6^ proteins co-enriched with endogenous FERRY subunits Fy1 or Fy2 relative to IgG controls following immunoprecipitation from M0 and LPS-stimulated macrophage lysates (n = 3 donors). Vertical dashed lines mark the fold-change cutoff (±log₂(1.5)) and the horizontal dashed line marks p = 0.05. Proteins exceeding both thresholds in the upper right quadrant are considered significantly enriched. Points are colored by protein category: FERRY complex subunits (light blue); RNA-binding proteins (RBP^7^; dark blue); ribosomal proteins (GO annotation; green); and all other proteins (grey). (**a**) Fy2 (PPP1R21) immunoprecipitation from LPS-stimulated macrophages, shown as log₂(LPS-Fy2/LPS-IgG). All five FERRY subunits (Fy1-Fy5) were significantly co-enriched. (**b**) Fy1 (TBCK) immunoprecipitation from resting M0 macrophages, shown as log₂(M0-Fy1/M0-IgG). Colored boxes in the upper corners show the number of significantly enriched proteins in each category. (**c**) Fy1 (TBCK) immunoprecipitation from LPS-stimulated macrophages, shown as log₂(LPS-Fy1/LPS-IgG). (**d**) Comparison of proteins co-enriched with Fy1 (TBCK) in LPS-stimulated versus resting macrophages, shown as log₂(LPS-Fy1/M0-Fy1). (**e**) Gene Ontology molecular function (GO:MF) enrichment analysis of proteins significantly enriched in endogenous Fy1 (TBCK) immunoprecipitations relative to IgG controls from M0 and LPS-stimulated macrophages. Enrichment was determined using Fisher’s exact test on the CRAPome-filtered^6^ enriched protein set. Each dot represents a GO:MF term; terms with FDR < 0.05 and n > 5 proteins are shown. (**f**,**g**) Venn diagrams comparing proteins significantly enriched in endogenous Fy1 (TBCK vs IgG) and Fy2 (PPP1R21 vs IgG) immunoprecipitations from macrophages with (**f**) the published GST-FERRY interactome from HEK293 cells^8^ (proteins annotated as "FERRY-complex enriched") and (**g**) the published Fy2 (PPP1R21) AP-MS interactome from HeLa cells (Wang et al^9^.; supplementary Table S12, filtered to high-confidence, >1 peptide and abundance > 5000). (**h**,**i**) Representative PLA images detecting interactions between Fy2 and Fy4 (**h**) and corresponding negative controls (**i**). PLA signals are shown in red, and cell membranes and nuclei are shown in green and blue, respectively.

## Supplementary Information

### **(A)** Supplementary Data Set Legends

#### Data S1

**Table S1-1:** Differential gene expression analysis was performed with DESeq2, using its median-of-ratios method to normalize RNA counts.
| # | Title | Description |
| --- | --- | --- |
| 1 | S1-1 Bulk RNA Sequencing Data | Differential gene expression analysis of bulk RNA-seq data from M0 and TLR/STING-stimulated macrophages (related to Figure 1 and Extended Data Fig. 1, 2). |
| 2 | S1-2 qPCR Primer Sequences | Sequences of primers for qPCR for validation of RNA-seq (related to Extended Data Fig. 2) |

**Table S1-1:** Differential gene expression analysis was performed with DESeq2, using its median-of-ratios method to normalize RNA counts. Genes with zero mapped reads were excluded from display. Each stimulus was compared against M0 (denoted M0_Low in the corresponding GEO submission). Normalized counts for each stimulus and donor are shown to the right of the differential expression statistics. Genes are the union across the seven comparisons. Genes are restricted to the primary assembly. Data are from n = 3 donors.

**Table S1-2:** Forward and reverse primers for validation of RNA-seq targets and housekeeping genes by qPCR.

#### Data S2

**Table S2-1:** TMT-exp data showing protein expression changes in macrophages stimulated with TLR/STING agonists compared to M0 macrophages. Data are from n = 3 donors.
| # | Title | Description |
| --- | --- | --- |
| 1 | S2-1 Unenriched proteomics | Whole proteome data from M0 and TLR/STING-stimulated macrophages (related to Figure 1 and Extended Data Fig. 1, 2) |
| 2 | S2-2 Bulk RNA vs WP | Whole proteome and transcriptome data from M0 and TLR/STING-stimulated macrophages (related to Figure 1 and Extended Data Fig. 1, 2) |
| 3 | S2-3 GSEA of PC loadings | Gene set enrichment analysis (GSEA) of whole proteome PCA loadings (related to Extended Data Fig. 1) |
| 4 | S2-4 GO-term enrichment | GO-term enrichment analysis of proteomic data (related to Figures 2 and 3, and Extended Data Fig. 2, 5, and 7) |
| 5 | S2-5 Reactivity changes | Cysteine reactivity changes identified from IA-DTB reactivity TMT-ABPP data (related to Figures 2, 3, and 4, and Extended Data Fig. 3, 4, 5, and 6). |
| 6 | S2-6 Phosphoproteomics | Expression-normalized phosphoproteomics data from M0 and LPS-stimulated macrophages (related to Figure 3 and Extended Data Fig. 5) |
| 7 | S2-7<br>Immunoprecipitation<br>MS | Immunoprecipitation-mass spectrometry (IP-MS) data (related to Figure 5 and Extended Data Fig. 7) |
| 8 | S2-8 Cross-omics Venn | Per-protein membership behind the cross-omics GO-term Venn diagrams (related to Figure 3 and Extended Data Fig. 5) |

**Table S2-1:** TMT-exp data showing protein expression changes in macrophages stimulated with TLR/STING agonists compared to M0 macrophages. Data are from n = 3 donors.

**Table S2-2:** TMT-exp and bulk RNA-seq data showing gene and protein expression changes in macrophages stimulated with TLR/STING agonists compared to M0 macrophages. TMT-exp and bulk RNA-seq are from n = 3 donors each (different donors for each assay).

**Table S2-3:** Pre-ranked GSEA of the protein loadings on each principal component of the whole proteome PCA, against the GO biological process gene sets of the 2026 MSigDB. A positive NES marks a term enriched at the positive end of that principal component. Only significant terms (FDR q-value < 0.01) are shown.

**Table S2-4:** GO-term enrichment analysis of whole proteome, reactivity, and phosphoproteomic data. Fisher’s exact test and the 2026 MSigDB gene sets were used for all analyses, and the background for each test is specified in the table. Only terms reaching significance (FDR < 0.05 and more than 5 proteins, except where a panel’s own cutoff is stricter) are listed, so a panel may plot a non-significant point that has no row here. ‘Relevant figure’ gives the panel in which each term is plotted. Protein counts for the Cross-omics rows are the same change sets the Venn diagrams use; see Table S2-8 for their per-protein membership.

**Table S2-5:** Proteins with identified cysteine reactivity changes in at least one condition (TLR1-2, TLR3, TLR4, TLR7, TLR8, STING) compared to M0 macrophages. Data are median from n = 3 donors. Per-condition columns list one value per residue in the ‘residue’ column, ‘|’-separated; ‘(C1;C2)’ marks a change that could not be attributed to a single residue and counts once. The cross-reference columns on the right describe only the cysteines that change: complex membership (CORUM, ComplexPortal), conservation depth (out of 102 organisms), AlphaFold solvent accessibility (pPSE; buried > 9, intermediate > 5) and secondary structure, AlphaMissense pathogenicity (pathogenic > 0.564, benign < 0.34), and pathogenic ClinVar variants within 15 residues.

**Table S2-6:** Phosphosite quantification in macrophages stimulated with LPS (TLR4) compared to M0 macrophages. Per-replicate channel ratios were normalized to protein expression by dividing each phosphosite’s channel ratio by that protein’s whole proteome fold change, so changes reflect phosphorylation rather than protein abundance; phosphosites on proteins without whole proteome coverage are not reported. Data are from n = 4 donors x 3 technical replicates, restricted to phosphosites quantified in at least 2 donors. Significance is an unpaired two-sample t-test of all TLR4 channels against all M0 channels; p-values are raw and are not corrected for multiple testing. ‘Regulation’ marks a phosphosite Significant Up or Significant Down at p < 0.05 and a fold change greater than 2.

**Table S2-7:** Immunoprecipitation-mass spectrometry showing proteins co-enriched with TBCK or PPP1R21 in LPS-stimulated macrophages (TLR4) or unstimulated (M0) macrophages. Data are from n = 3 donors (different donors for each immunoprecipitation assay).

**Table S2-8:** One row per protein in the union of the three TLR4-vs-M0 change sets for each GO term, with a 1/0 flag per profiling method. Unlike Table S2-4, these are raw overlaps of the term’s gene list with each method’s significant changes, not an enrichment test, so no p-values apply; summing a flag column gives that method’s circle total in the panel. ‘GO term source’ gives the gene set used, which for GTPase activity is a union of GTPase GO terms rather than GO:0003924 alone.

#### Data S3

**Table S3-1:** Polar metabolite abundance values in TLR4-stimulated macrophages compared to M0 macrophages as determined by LC-MS/MS analysis.
| # | Title | Description |
| --- | --- | --- |
| 1 | S3-1 Polar metabolites | Abundance of polar metabolites in M0 and TLR4-stimulated macrophages (related to Figure 2 and Extended Data Fig. 3) |

**Table S3-1:** Polar metabolite abundance values in TLR4-stimulated macrophages compared to M0 macrophages as determined by LC-MS/MS analysis. The channel ratio values after quantile regression imputation of left censored data (QRILC) are shown alongside raw signal intensity values and differential abundance calculations. Data are from n = 4 donors.

#### Data S4

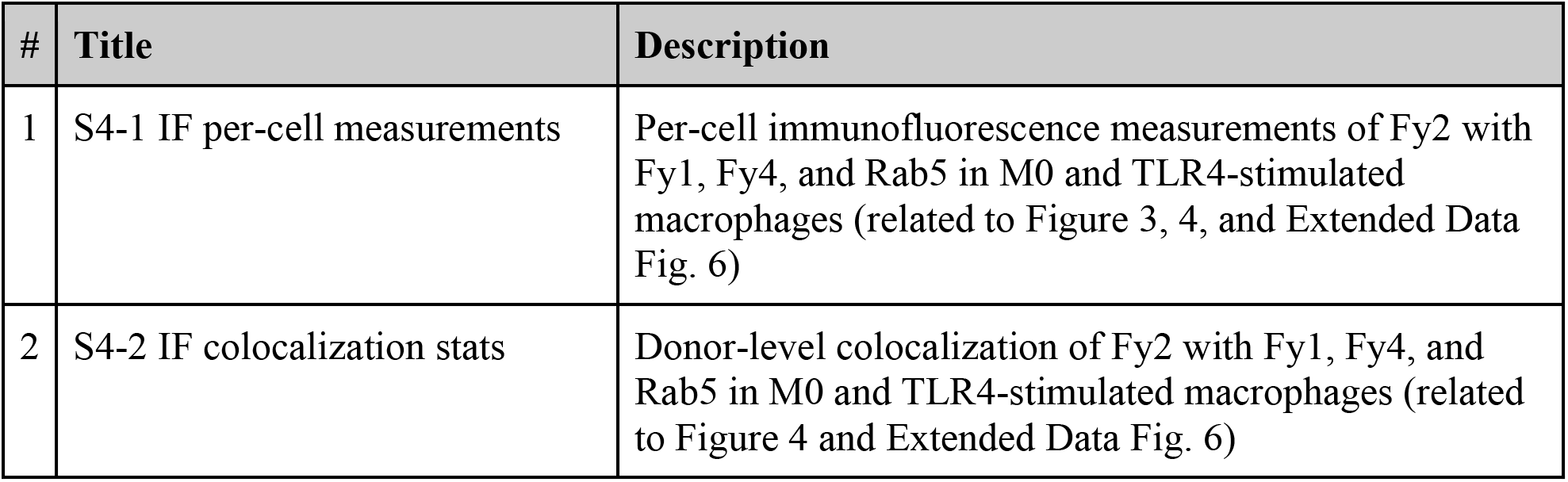

**Table S4-1:** Per-cell vesicle counts, sizes, and intensities, and split colocalization coefficients of Fy2 with Fy1, Fy4, or Rab5 in M0 and TLR4-stimulated macrophages, one row per cell.
| # | Title | Description |
| --- | --- | --- |
| 1 | S4-1 IF per-cell measurements | Per-cell immunofluorescence measurements of Fy2 with Fy1, Fy4, and Rab5 in M0 and TLR4-stimulated macrophages (related to Figure 3, 4, and Extended Data Fig. 6) |
| 2 | S4-2 IF colocalization stats | Donor-level colocalization of Fy2 with Fy1, Fy4, and Rab5 in M0 and TLR4-stimulated macrophages (related to Figure 4 and Extended Data Fig. 6) |

**Table S4-1:** Per-cell vesicle counts, sizes, and intensities, and split colocalization coefficients of Fy2 with Fy1, Fy4, or Rab5 in M0 and TLR4-stimulated macrophages, one row per cell. In column headers, the partner of Fy2 in each antibody pair is referred to as “marker”. Colocalization coefficients are shown in both directions: Coloc. (X/Y) is the fraction of the X signal on Y. Puncta with a radius < 0.15 µm were excluded. No chromatic aberration correction was applied. Data are from n = 6 donors.

**Table S4-2:** Donor-level split colocalization coefficients for each direction of each antibody pair (intensity-weighted). Per-cell values were averaged per donor and condition, and the mean ± SEM of donor means is shown; P values are from a two-tailed paired t-test on the donor means. The number of cells quantified per condition is listed for each pair. Data are from n = 6 donors.

### (B) Biological Methods

#### EXPERIMENTAL MODEL AND SUBJECT DETAILS

##### Human samples

All studies involving primary human macrophages were performed using buffy coats obtained from the New York Blood Center, leukopaks and LRS cones from STEMCELL Technologies, and blood and leukapheresis samples from the Rockefeller University Hospital. Blood samples were collected from randomly selected, de-identified healthy donors with donor sex not recorded. The human subjects protocol was reviewed and approved annually by the Rockefeller University Hospital Institutional Review Board (IRB; protocol EVI-1024).

##### Isolation of human peripheral blood mononuclear cells (PBMCs) and monocytes

Peripheral blood was diluted with PBS, and 25 mL of the diluted cells were layered over 12.5 mL of Lymphoprep (STEMCELL Technologies) using 50 mL Falcon tubes. Tubes were centrifuged at room temperature (900 x *g*, 25 °C, 20 min) with the acceleration set to 50% and brake disabled. Following the spin, the plasma and cell layers were transferred to new 50 mL Falcon tubes, diluted with PBS, and pelleted (520 x *g*, 25 °C, 8 min). The pellet was washed once in PBS, resuspended in 5 mL of 1X RBC lysis buffer (Santa Cruz Biotechnologies), and incubated for 5 min at room temperature. Cells were then diluted with PBS, pelleted (520 x *g*, 25 °C, 8 min), and washed with PBS. PBMCs were resuspended at 50 × 10⁶ cells /mL, and monocytes were isolated by negative selection using the EasySep Monocyte Enrichment Kit without CD16 Depletion (STEMCELL Technologies) according to manufacturer’s instructions.

#### METHODS DETAILS

##### Macrophage isolation and polarization

Freshly isolated monocytes were resuspended at 1 × 10⁶ cells/mL in macrophage media, consisting of RPMI-1640 supplemented with 10% FBS (Omega Scientific), 2 mM GlutaMAX (Gibco), 100 U/mL Penicillin-Streptomycin (Gibco), 1 mM Sodium Pyruvate (Gibco), 2 mM HEPES pH 7.0 (Boston Bioproducts), and 50 ng/mL mCSF (STEMCELL Technologies). Cells were seeded at 15 × 10⁶ cells per 15 cm culture plate, 5 × 10⁶ cells per 10 cm culture plate, or 0.83 × 10⁶ cells per well of a 6-well plate and cultured at 37 °C in a humidified 5% CO_2_ incubator for three days. Cells were washed three times with PBS and re-supplemented with macrophage media containing mCSF for another three days. Cells were then washed three times with PBS and re-supplemented with macrophage media containing mCSF and TLR or STING agonists (see table below) for 24 h. Cells were then placed on ice, washed 3x with ice-cold PBS, and harvested into ice-cold PBS by scraping. Cells were then pelleted (520 x *g*, 4 °C, 8 min), flash frozen, and kept at -80 °C until further analysis.

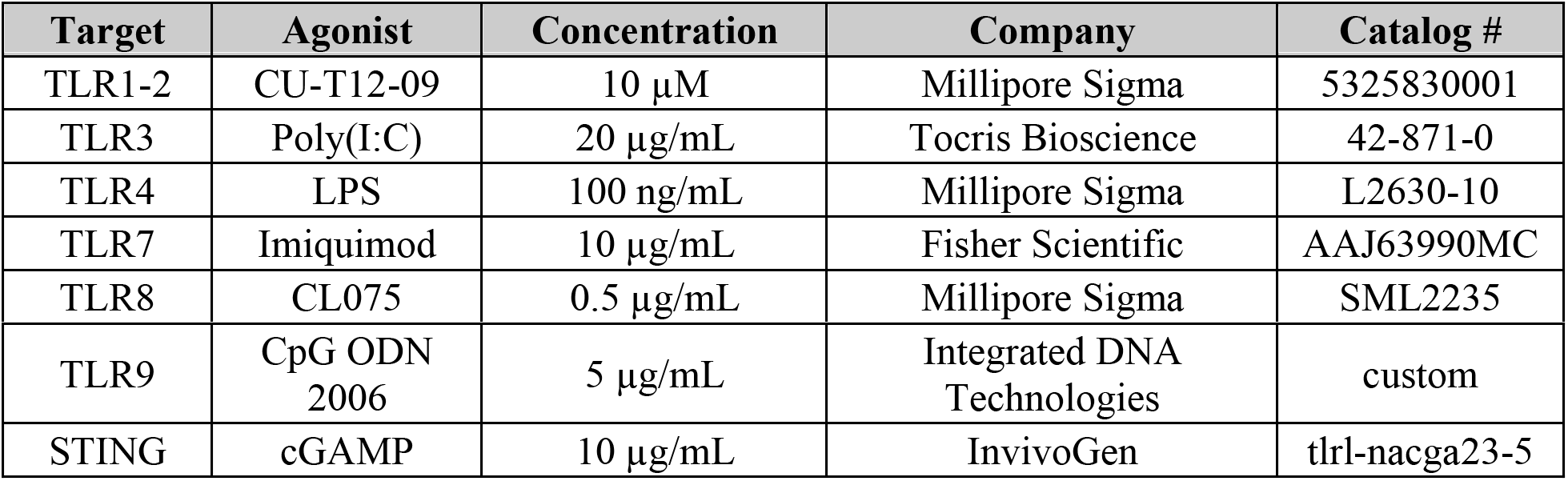

##### Flow cytometry analysis

Polarized cells were washed once with room temperature PBS and incubated with TrypLE (Gibco, 12604021) for 30 min at 37 °C in a humidified 5% CO_2_ incubator, with vigorous tapping on the edges of the plate every 10 min. Cells were then gently scraped and transferred into a tube. The plate was then washed with PBS, and the wash was pooled with previously collected cells. Cells were spun down (520 x *g*, 4 °C, 8 min), resuspended in PBS, and plated at 100,000 live cells per well in a U-bottom 96-well plate. Cells were spun down (520 x *g*, 4 °C, 5 min) and washed once with PBS. Cells were then resuspended in Live/Dead Fixable NIR Viability Dye (Invitrogen, L10119) at a 1:50,000 dilution in PBS, and incubated at room temperature for 15 min in the dark. Cells were then spun down and washed once with PBS. Cells were then incubated with 25 µL of Fc Block (BioLegend, 422302) and Monocyte Block (BioLegend, 426102) both at a 1:20 dilution in staining buffer (1% FBS in PBS) for 15 min at 4 °C. Cells were then stained for surface markers at a 1:400 dilution in Brilliant Stain buffer and incubated on ice for 25 min at 4 °C. Cells were then diluted with 150 µL of staining buffer, spun down, and washed once more with staining buffer. Cells were then resuspended in 4% PFA (FUJIFILM) for 15 min at 4 °C. Cells were spun down, washed once with staining buffer, and resuspended in staining buffer with 2 mM EDTA. Data was acquired using an Agilent Novocyte Penteon cytometer, with a minimum of 4,000 live cells measured per condition. Data was reported as geometric mean of individual markers with a gate set using fluorescence minus one (FMO) controls and normalized to the M0 condition. Significance was calculated by a matched one-way ANOVA with Dunnett’s multiple comparisons test (GraphPad Prism, Version 10).

| Staining type | Marker | Fluorophore | Catalog # | Clone | Dilution |
| --- | --- | --- | --- | --- | --- |
| Surface | CD14 | BV650 | 563419 | M5E2 | 1:400 |
| Surface | CD11b | BV711 | 563168 | M1/70 | 1:400 |
| Surface | CD80 | PE | 560925 | L307.4 | 1:400 |
| Viability | Viability | Live/Dead NIR | L10119 | -- | 1:50,000 |

##### Enzyme-Linked Immunosorbent Assay (ELISA)

Following the 24 h incubation with TLR or STING agonists, macrophage media was taken from cells into a new 96-well plate, spun briefly (520 x g, 4 °C, 5 min) to remove cell debris, and the supernatant containing cytokines was stored at -80 °C until analysis. ELISA was performed with IL-6 or TNFα DuoSet ELISA Kits (R&D Systems, DY206 and DY210) according to manufacturer’s instructions. Absorbance was read out on a plate reader at both 450 nm with a 540 nm background subtraction, and concentrations were interpolated from standards fit to a 4-PL model. Significance was calculated by a matched one-way ANOVA with Dunnett’s multiple comparisons test (GraphPad Prism, Version 10) on log_2_-transformed concentration values.

##### RNA isolation for qPCR and RNA sequencing analysis of basal state and activated macrophages

Following stimulation, plates containing cells were put on ice, washed three times with ice-cold PBS, and the cells were harvested by scraping into ice-cold PBS and transferring to new Falcon tubes. Cells were then pelleted (520 x g, 4 °C, 5 min), transferred to new low binding Eppendorf tubes, resuspended in TRIzol reagent (Invitrogen, 15-596-026), snap-frozen in liquid nitrogen, and stored at -80 °C. RNA was extracted using the Direct-zol RNA Miniprep kit (Zymo Research, R2052) with on-column DNase I digestion according to manufacturer’s instructions. Purified RNA was flash frozen and stored at -80 °C until further analysis.

##### RNA sequencing

###### Sample preparation

RNA quality and concentration were assessed using an Agilent Bioanalyzer, and samples with a RIN > 8 were used for the library prep. 100 ng of total RNA was used to generate RNA-seq libraries with the Illumina stranded mRNA kit (20040534) according to the manufacturer’s protocol. Libraries were prepared with unique dual indexes and pooled at equal molar ratios. The pool was denatured and sequenced on an Illumina NovaSeq 6000 sequencer using V1.5 reagents, S2 flowcell, and NovaSeq Control Software V1.7.0 to generate 2 x 100 bp paired-end reads, according to the manufacturer’s protocol.

###### Data processing

Paired-end RNA-seq reads were aligned to the GRCh38 reference genome using STAR (v2.7.7a) in two-pass mode. The STAR genome index was generated using GENCODE release 44 (Ensembl 110) gene annotations. Gene-level counts were generated with featureCounts (v1.4.6-p5) in paired-end mode using the corresponding GENCODE release 44 annotations.

###### Principal Component Analysis

The 5,000 protein-coding genes with the highest row-wise variance were used and normalized count values were log_2_-transformed before principal component analysis (PCA). PCA was performed using *scikit-learn* library for Python. The first two principal components (PC1 and PC2) were plotted, and loadings were visualized for the five genes with the highest and lowest loadings along each component.

##### RT-qPCR

RNA concentration was quantified using a Qubit fluorometer with the Qubit RNA Broad Range (BR) kit (Invitrogen, Q10210). cDNA was synthesized using the iScript cDNA Synthesis Kit (Bio-Rad, 1708891) according to manufacturer’s instructions. qPCR was performed on a QuantStudio5 (Thermo Fisher) in standard cycling mode with a terminal melt curve step in a 384-well plate. Each reaction contained 50 ng of cDNA input, 400 nM of each primer, and 1X Select Master Mix (Applied Biosystems, 4472908) at a final volume of 10 µL per well. Primer sequences are listed in Supplementary Table S1-2. Samples were run in triplicate and quantified by the standard curve method. No-template (NTC) and no-reverse transcriptase (-RT) controls were included on each plate. Gene expression was normalized to the average of housekeeping genes ACTB and C1orf43, which were chosen due to their unchanging expression in our RNA sequencing data, and expressed as log_2_(fold change) relative to unstimulated (M0) control conditions. Statistical comparisons were performed by one-way ANOVA with post-hoc multiple comparisons testing (GraphPad Prism, Version 10)

##### Metabolomic analysis

###### Sample preparation

Monocytes were plated on 10 cm plates and differentiated as described above. Macrophages were then polarized with either fresh macrophage media or media supplemented with agonists for 24 h. Plates were then transferred to ice and washed twice with ice-cold PBS. Cells were scraped directly into 1 mL of ice-cold 80% MeOH in water and transferred to an Eppendorf tube. Cells were then vortexed for 45 s and incubated on dry ice for 5 min. Cells were again vortexed for 45 s and pelleted to get rid of debris (14,000 x *g*, 4 °C, 20 min). Supernatant was transferred to a clean tube and dried under flow of Nitrogen gas. Samples were stored at -80 °C until analysis.

###### Polar metabolite analysis

The sample was redissolved in HPLC grade water before it was analyzed by LC-MS/MS. Metabolites were measured on an Orbitrap Exploris 240 mass spectrometer (Thermo Scientific) coupled to a Vanquish UPLC system (Thermo Scientific). A Sequant ZIC-pHILIC column (2.1 mm i.d. × 150 mm, particle size of 5 µm, Millipore Sigma) was used for separation of metabolites. A 2.1 × 20 mm guard column with the same packing material was used for protection of the analytical column. Flow rate was set at 150 μL/min. Buffers consisted of 100% acetonitrile for mobile phase A, and 0.1% NH_4_OH/20 mM CH_3_COONH_4_ in water for mobile phase B. The chromatographic gradient ran from 85% to 30% A in 20 min followed by a wash with 30% A and re-equilibration at 85% A. The Orbitrap Exploris 240 was operated in full scan, polarity-switching mode with the following parameters: the spray voltage is 3.0 kV in positive mode and 2.5 kV in negative mode, the heated capillary temperature 275 °C, the HESI probe temperature 320 °C, the sheath gas flow 35 units, the auxiliary gas flow 7 units. MS data acquisition was performed in the m/z range of 70–1,050, with 90,000 resolution (at 200 m/z). The AGC target was 1e6 and the maximum injection time was 250 ms. The MS data was processed using XCalibur 4.1 (Thermo Scientific) to obtain the metabolite signal intensities. Identification required exact mass (within 5 ppm) and standard retention times.

###### Data processing and imputation

Within each biological replicate, metabolites were required to have at least one condition with an average signal intensity above 5,000 (179/205 metabolites passed). After filtering and log transformation, missing data was imputed through quantile regression imputation of left-censored data (QRILC) performed using imputeLCMD version 2.1. Channel ratios (signal intensity / sum of signal intensities per metabolite) were calculated and used for all analyses.

###### Hierarchical clustering

The top 25 increasing and decreasing metabolites by p-value (T-test between M0/TLR4) were used for hierarchical clustering. A z-score was calculated for each observation by z = (x-μ)/σ, where x is the channel ratio, μ is the mean channel ratio of the metabolite across all replicates, and σ is the standard deviation of the channel ratio of the metabolite across all replicates. Hierarchical clustering was performed using ComplexHeatmap^1^ library version 2.10.0 for R version 4.1.1. The distance matrix was calculated using the “euclidean” method and clustering was performed using the “complete” method. Metabolite class annotations were curated through literature review.

##### Proteomic platforms: Abundance-based proteomics with TMT multiplexing

###### Sample preparation

Three 16-plex experiments were performed with primary human macrophages from different donors, consisting of technical duplicates from 8 different macrophage states. Frozen cell pellets were thawed on ice and resuspended in PBS with protease inhibitor cocktail (Roche cOmplete Mini, EDTA-free, 1 tablet per 10 mL PBS). Cells were then lysed with sonication (three cycles of 8 x 1s pulses on a 60% duty cycle). Protein concentration was determined with the DC assay (Bio-Rad), and lysates were normalized to 2 mg/mL.

For each sample, 100 µL (200 µg) of lysate was added to 48 mg urea to a final concentration of 8 M. Freshly made stock of DTT (5 µL, 200 mM in water) was added to each sample and incubated at 65 °C for 15 min. Freshly made stock of iodoacetamide (5 µL, 400 mM in water) was then added and incubated for 30 min at 37 °C in the dark with shaking. Proteins were precipitated by addition of 500 µL ice-cold methanol, 400 µL water, and 150 µL chloroform, followed by centrifugation (10,000 x *g*, 4 °C, 10 min). The upper aqueous layer was carefully removed without disturbing the protein disc, and an additional 400 µL of cold methanol was added. Pellets were resuspended by sonication and centrifuged (10,000 x *g*, 4 °C, 10 min). The supernatant was discarded, and the pellets were resuspended by sonication in 200 mM EPPS buffer (pH 8.0). The proteins were then digested with 2.5 µg of Trypsin/LysC (Promega) with CaCl_2_ (1 mM final concentration) overnight at 37 °C with shaking.

###### TMT labeling

Digests were centrifuged at room temperature (10,000 x *g*, 25 °C, 5 min) to remove undigested material and supernatants were transferred to new tubes. Peptide concentration was measured with the microBCA assay (Thermo Fisher, 23235). For each sample, 25 µg of peptides were then transferred to a new tube, topped up to equal volumes, and acetonitrile was added to each tube to a final concentration of 25%. Samples were then labeled with 100 µg of corresponding TMT tags (Thermo Fisher, A44521; 20 µg/µL in acetonitrile) for 1 h at room temperature. Labeling reactions were quenched with 6 µL of 5% Hydroxylamine in water (Sigma-Aldrich, 467804) for 15 min at room temperature and acidified with formic acid (VWR, 94318) to a final concentration of 5%.

###### Normalization of TMT input and desalting

A ratio check sample was prepared by combining a small aliquot from each labeled sample, desalting using a C18 stage tip (Thermo Fisher), and drying it in a SpeedVac. The sample was resuspended in Buffer A (5% acetonitrile, 95% water, 0.1% formic acid) and analyzed using an Orbitrap Eclipse mass spectrometer using a 70-min LC gradient: 0-10 min, 5% buffer B (0.1% FA in acetonitrile) in buffer A (0.1% FA in water); 10-50 min, linear increase from 5% to 20% buffer B; 50-55 min, ramp to 45% buffer B; 55-57 min, ramp to 95% buffer B; 57-59 min, hold at 95% buffer B; 59-61 min, decrease to 5% buffer B; 61-63 min, ramp back to 95% buffer B; 63-70 min: re-equilibration at 5% buffer B. Normalization factors (N) were calculated for each TMT channel, and volume equivalents of 10 µL x N per channel were pooled and dried in the SpeedVac. The pooled sample was then resuspended in Buffer A and acidified with formic acid to a final pH of < 3. A C18 SepPak cartridge (Waters, WAT054955) was activated with 3 x 1 mL of acetonitrile and equilibrated with 3 x 1 mL of Buffer A. Acidified sample was added to the equilibrated column, and the flow-through was reloaded for maximum enrichment. The column was then washed three times with Buffer A, eluted in 1 mL of Buffer B (80% acetonitrile, 20% water, 0.1% formic acid), and dried on the SpeedVac.

###### High pH HPLC fractionation (general protocol for all LC-MS/MS/MS workflows)

High pH HPLC fractionation was performed following previously published protocols^2^. Briefly, the desalted peptides were resuspended in 500 µL of Buffer A and fractionated offline using an Agilent HPLC system into a 96-well plate containing 10 µL of 20% formic acid. Separation was performed on a Zorbax Extend-C18 column (3.5 µm, 4.6 × 250 mm) at a flow rate of 0.5 mL/min using a 10 mM ammonium bicarbonate (Buffer C) to acetonitrile (Buffer D) gradient over 125 min. The gradient is as follows: 0-2 min, 100% Buffer C; 2-3 min, 0% to 13% Buffer D; 3-60 min, 13% to 50% Buffer D; 60-70 min, 50% to 80% Buffer D; 71-75 min, 100% Buffer D; 75-76 min, 100% to 0% Buffer D; 76-85 min, 100% Buffer C; 85-88 min, 0% to 13% Buffer D; 88-90 min, 13% to 80% Buffer D; 90-95 min, hold at 80% Buffer D; 96-101 min, 100% Buffer C; 101-104 min, 0% to 13% Buffer D; 104-106 min, 13% to 80% Buffer D; 106-111 min, hold at 80% Buffer D; 111-112 min, 80% to 0% Buffer D. After fractionation, the plate was dried in a SpeedVac. The peptides in each well were then resuspended in Buffer B, and the fractions were consolidated into 12 final fractions by combining wells in each column of the plate. Fractions were dried in the SpeedVac and used for LC-MS/MS/MS analysis.

###### General protocol for liquid chromatography mass spectrometry analysis of TMT-multiplexed samples

TMT-labeled peptides were analyzed by LC-MS/MS/MS using a 160-min gradient on an Orbitrap Eclipse mass spectrometer (Thermo Fisher) as described before^2, 3^. MS1 spectra were acquired in the Orbitrap at a resolution of 120,000 over a mass range of 400-1600 m/z, with a 40% RF lens setting, automatic gain control (AGC) target set to 250% (normalized), and maximum injection time set to automatic. Dynamic exclusion was enabled with a repeat count of 1 and an exclusion duration of 60 s. The ten most abundant precursor ions from the MS1 scan were selected for MS2/MS3 analysis, isolated using the quadrupole with a 0.7 m/z isolation window, and fragmented by collision-induced dissociation (CID) in the ion trap (standard AGC, 35% normalized collision energy (NCE), and a 120 ms maximum injection time). Synchronous precursor selection (SPS) was enabled to select up to 10 MS2 fragment ions for MS3 analysis. MS3 spectra were acquired in the Orbitrap following high-energy collision-induced dissociation (HCD) with 55% NCE, a normalized AGC target of 500%, a maximum injection time of 118 ms, a resolution of 60,000, and an 0.7 m/z isolation window for MS3.

Raw data files were processed using RAW Converter (v1.1.0.22; available at github.com/proteomicsyates/RawConverter), and the resulting MS1, MS2, and MS3 files were uploaded to the Integrated Proteomics Pipeline (IP2). Peptide identification was performed using the ProLuCID algorithm against a reverse concatenated, non-redundant version of the Human UniProt database (release 2024_03). Cysteine residues were searched with a static modification for carbamidomethylation (+57.02146 Da). A differential modification for iodoacetamide desthiobiotin labeling (+398.25292 Da) was included for site-of-labeling experiments. TMT labeling was considered as a static modification on peptide *N-*termini and lysine residues (+229.1629 Da for TMT 10-plex, +304.2071 Da for TMT 16-plex). Search results were filtered using DTASelect (v2.0) to maintain a peptide-level false discovery rate (FDR) below 1%. Quantification was performed using MS3 reporter ions with a mass tolerance of 20 ppm via the IP2 platform.

##### Data processing and analysis for TMT-exp experiments

###### Data processing

Three 16-plex experiments with primary human macrophages from different biological donors were processed. Proteins were required to have at least two unique quantified peptides to pass into the final list. Peptide ratios (signal intensity/sum of signal intensities per peptide) were calculated and ratios of all peptides per protein were averaged. Keratins and contaminants were not included in analysis. Proteins needed to be quantified in at least 2 experiments to be included in the final table.

###### Principal Component Analysis

Principal component analysis was performed in Python using the scikit-learn library^4^. Protein ratio values were log₂-transformed, and the analysis was restricted to proteins quantified across all experiments. The first two principal components (PC1 and PC2) were plotted, and loadings were visualized for the five proteins with the highest and lowest loadings along each component.

###### Gene set enrichment analysis (GSEA) of PCA loadings

PCA loadings were used as the ranking metric for GSEA, performed with the pre-rank function in GSEApy v1.1.9 against the Gene Ontology biological process database ("c5.go.bp"). The top 15 enriched terms, ranked by normalized enrichment score (NES), were plotted.

###### Protein half-life analysis

Protein half-lives were referenced from the monocyte turnover measurements in Kosiński *et al^5^.* Each protein’s half-life was defined as the mean of two biological replicates (or the single available value), and mapped to the macrophage differential-expression values by gene symbol. Half-lives were binned (0-24, 24-48, 48-96, 96+ h) and their distribution was analyzed based on the observed concordance between transcript- and protein-abundance changes for both upregulated and downregulated proteins separately.

###### Hierarchical clustering

Whole-proteome TMT signals were calculated and used as log_2_ fold changes relative to the mean of the M0 channels, and gene-set proteins quantified across all replicates were displayed with ComplexHeatmap. Rows were ordered by hierarchical clustering (Euclidean distance, complete linkage); columns were grouped by agonist condition.

##### Proteomic platforms: Cysteine reactivity profiling, TMT-ABPP

###### Sample preparation

Three 16-plex experiments were performed with primary human macrophages from different donors, consisting of technical duplicates from 8 different macrophage states. Frozen cell pellets were thawed on ice and resuspended in PBS. Cells were then lysed with sonication (three cycles of 8 x 1 s pulses on a 60% duty cycle). Protein concentration was determined with the DC assay, and lysates were normalized to 2 mg/mL.

For each sample, 500 µL (1 mg) of lysate was transferred into low-bind Eppendorf tubes and labeled with 5 µL of 10 mM iodoacetamide-desthiobiotin probe (IA-DTB) for 1 h at room temperature. Proteins were then precipitated by adding 500 µL ice-cold methanol and 100 µL chloroform to each tube, vortexing, and centrifuging (10,000 x *g*, 10 min, 4 °C). The supernatant was removed without disturbing the protein disc, and an additional 400 µL of cold methanol was then added to each sample. The pellets were resuspended by sonication, centrifuged (10,000 x *g*, 10 min, 4 °C), and resuspended by sonication in 90 µL of 50 mM TEAB buffer (pH 8.5) containing 8 M urea and 10 mM DTT. Samples were then heated at 65 °C for 20 min, followed by addition of freshly prepared 10 µL of 500 mM iodoacetamide in water and incubation for 30 min at 37 °C in the dark with shaking.

###### Trypsin/LysC digestion and streptavidin enrichment

Samples were diluted with 300 µL of 50 mM TEAB, and proteins were digested overnight at 37 °C with shaking using 2 µg of Trypsin/LysC and 4 µL of 100 mM CaCl_2_. The next day, Streptavidin Agarose Beads (50 µL per sample, Pierce, 20353) were washed twice with Wash Buffer 1 (WB1: 50 mM TEAB, 150 mM NaCl, 0.2% NP-40), resuspended in 300 µL WB1 per sample, and added to the digested samples. The samples were rotated at room temperature for 3 h. After enrichment, the samples were transferred to BioSpin columns (Bio-rad), washed three times each with Wash Buffer 2 (WB2: 50 mM TEAB, 150 mM NaCl, 0.1% NP-40), PBS, and water. The peptides were eluted in 50% acetonitrile : water with 0.1% formic acid and dried in the SpeedVac.

###### TMT labeling

The enriched peptides were resuspended in 100 µL of 30% acetonitrile in 200 mM EPPS (pH 8.0) and labeled with 3 µL (20 µg/µL in acetonitrile) TMT reagent for 1 h at room temperature, as described above. Reactions were quenched with 3 µL of 5% Hydroxylamine for 15 min at room temperature and acidified with 5 µL of formic acid. Labeled peptides from all samples were then pooled and dried in the SpeedVac. Desalting, high-pH fractionation, and LC-MS/MS/MS analysis were performed as described above, with the IA-DTB modification (+398.25292 Da) included as a variable Cysteine modification during the searches.

##### Data processing and analysis for TMT-ABPP Data quality filters

Peptides were required to be quantified in at least two biological replicates, and proteins to have at least two quantified peptides. Technical replicates were averaged before filtering. A percent-control value was discarded when the mean signal intensity of its channel was below 5,000 and the corresponding M0 (control) channel was also below 5,000. Residues quantified in exactly two replicates were excluded from analysis if at least one replicate showed less than a 1.5-fold change and the two replicates differed from each other by 2-fold or more. Proteins manually annotated as low quality were excluded.

##### Reactivity-change criteria

A cysteine was called a reactivity change when all of the following held. Throughout, a cysteine’s "ratio to another cysteine" means its largest fold-difference from other cysteine quantified on the same protein in that condition.

1. At least one cysteine on the protein fell within 1.5-fold of the protein’s expression value, or the protein carried cysteines both more than 1.5-fold above and more than 1.5-fold below it. A cysteine whose own ratio to another cysteine exceeded 3-fold satisfied this criterion regardless.
2. At least one cysteine on the protein changed less than 2-fold, or the cysteine’s ratio to another cysteine exceeded 3-fold.
3. The cysteine’s ratio to another cysteine exceeded 2-fold in at least two replicates. Where a peptide was missing from a replicate, its mean value across the remaining replicates was substituted.
4. Where whole-proteome expression data were available, the cysteine’s ratio to the protein’s expression value exceeded 2-fold.
5. For proteins with two to four peptides, the cysteine’s ratio to another cysteine exceeded 2-fold. For proteins with five or more peptides – and only when at least one cysteine on the protein changed less than 2-fold – the cysteine instead differed by more than 2-fold from the protein’s median cysteine value. Where expression data were available, this difference was required to be in the same direction as the cysteine’s ratio to expression.

Cysteine reactivity changes on proteins with fewer than five peptides and no expression data could not be attributed to an individual residue; these were reported as ambiguous groups (Cys1;Cys2) and counted as a single reactivity change.

##### Proteomic platforms: Phosphoproteomics

###### Sample preparation

Four 6-plex experiments were performed with primary human macrophages from different donors, consisting of technical triplicates from M0 and TLR4-stimulated macrophages. Frozen cell pellets were thawed on ice and resuspended in 8 M urea lysis buffer (8 M urea in 10 mL of 200 mM EPPS pH 8.5 with 1 tablet each of Roche cOmplete Mini, EDTA-free protease inhibitor and PhosSTOP (Roche, 4906845001). Cells were then lysed with sonication (three cycles of 30 x 1 s pulses on a 80% duty cycle). Samples were then passed 10 times through a 25G needle. Protein concentration was determined with the BCA assay, and lysates were normalized to 3.7 mg/mL.

For each sample, 540 µL (2 mg) of lysate was transferred into 2 mL low-bind Eppendorf tubes and reduced with 20 µL of 300 mM DTT at 65 °C for 15 min. Proteins were then alkylated with 40 µL of 750 mM iodoacetamide for 30 min at 37 °C. Proteins were then precipitated by adding 600 µL ice-cold methanol and 120 µL chloroform to each tube, vortexing, and centrifuging (10,000 x *g*, 10 min, 4 °C). The supernatant was removed without disturbing the protein disc, and an additional 400 µL of cold methanol was then added to each sample. The pellets were resuspended by sonication, centrifuged (10,000 x *g*, 10 min, 4 °C), and resuspended by sonication in 800 µL of 200 mM EPPS (pH 8.5). Samples were digested with 5 µg of Trypsin/LysC resuspended at 0.5 µg/µL in Resuspension Buffer (Promega) in the presence of 8 µL 100 mM CaCl_2_ at 37 °C overnight with shaking.

The next day, samples were acidified with Formic Acid until the pH was below 3. Samples were spun down (10,000 x *g*, 1 min, 25 °C) to pellet undigested precipitates, and supernatant was transferred to a new low-bind tube. C18 SepPak cartridges were conditioned with acetonitrile (3 x 1 mL) and equilibrated with Buffer A (3 x 1 mL). Samples were loaded into individual cartridges, and flow-through was loaded into the cartridge again for maximum peptide binding. Cartridges were then washed with Buffer A (3 x 1 mL) and eluted in 1 mL of Buffer B. Samples were then dried in the SpeedVac.

###### TiO_2_ Enrichment

Samples were enriched with the High Select TiO_2_ Phosphopeptide Enrichment Kit (Thermo Scientific, A32993) according to manufacturer’s protocol. Briefly, all samples were resuspended in room temperature Binding/Equilibration Buffer. TiO_2_ spin tips were washed with Wash Buffer (2 x 20 µL), Binding/Equilibration Buffer (2 x 20 µL), and samples were loaded twice onto the tips. Tips were then washed again with Binding/Equilibration Buffer (2 x 20 µL), Wash Buffer (2 x 20 µL), Binding/Equilibration Buffer (2 x 20 µL), Wash Buffer (2 x 20 µL), and water (1 x 20 µL). Samples were eluted twice with 50 µL Elution Buffer and dried in the SpeedVac. All flow-through and washes were saved and pooled per sample and dried in the SpeedVac.

###### Fe-NTA Enrichment

Flow-through and washes were enriched with the High Select Fe-NTA Phosphopeptide Enrichment Kit (Thermo Fisher, A32992) according to manufacturer’s protocol. Briefly, spin columns were washed in Binding/Wash Buffer (2 x 200 µL) and the bottoms were capped with a Luer plug. Samples were resuspended in Binding/Wash Buffer and added to the column and incubated for 30 min at room temperature, with tapping every 10 minutes to resuspend resin. After incubation, samples were allowed to flow through the column, and columns were then washed with Binding/Wash Buffer (3 x 200 µL), LC-MS grade water (2 x 200 µL), and eluted twice with 100 µL of elution buffer. Samples were dried immediately in the SpeedVac.

###### TMT Labeling

Elutions from both Fe-NTA and TiO_2_ enrichments were resuspended in 100 µL 30% ACN in 200 mM EPPS (pH 8.0), vortexed into suspension, and spun briefly (500 x *g*, 10 s, RT). 5 µL of TMT tag at 20 µg/µL was added to each channel and incubated at room temperature for 1 h. After incubation, 5 µL of 5% hydroxylamine was added to each sample to quench the reaction and incubated for 15 min at room temperature. The sample was then acidified in 8 µL of Formic Acid. Samples were then pooled and dried in the SpeedVac, with Fe-NTA and TiO_2_ enrichments being kept separate. Desalting and high-pH fractionation were performed as described above.

###### General protocol for liquid chromatography mass spectrometry analysis of phosphoproteomics samples

TMT-labeled peptides were analyzed by LC-MS/MS/MS using a 160-min gradient on an Orbitrap Eclipse mass spectrometer (Thermo Fisher). MS1 spectra were acquired in the Orbitrap at a resolution of 120,000 over a mass range of 400-1500 m/z, with a 30% RF lens setting, automatic gain control (AGC) target set to Standard, and maximum injection time set to automatic. Monoisotopic peak determination was set to “Peptide”, with the isolation window centered on the most abundant peak. Candidate precursors were required to exceed an intensity threshold of 5.0 x 10^4^ and a 50% fit threshold within a 0.7 m/z fit window, and carry a charge state of 2-6. Dynamic exclusion was enabled with a repeat count of 1 and an exclusion duration of 60 s (± 10 ppm mass tolerance). The ten most abundant precursor ions from the MS1 scan were selected for MS2/MS3 analysis, isolated using the quadrupole with a 0.7 m/z isolation window and fragmented by collision-induced dissociation (CID) in the ion trap (standard AGC, 30% normalized collision energy (NCE), a 10 ms activation time, activation Q of 0.25, and a 54 ms maximum injection time). Multistage activation was enabled with a neutral loss mass of 97.9673, and resulting fragment spectra were detected in the Orbitrap at a resolution of 30,000. Synchronous precursor selection (SPS) was enabled to select up to 10 MS2 fragment ions for MS3 analysis. MS3 spectra were acquired in the Orbitrap following high-energy collision-induced dissociation (HCD) with 65% NCE, a normalized AGC target of 200%, a maximum injection time of 200 ms, a resolution of 50,000, and an 0.7 m/z isolation window for MS3.

###### Data analysis and processing

Differential phosphorylation of serine, threonine, and tyrosine (S/T/Y) was searched allowing a maximum of 1, 2, 3, and 10 differential modifications per peptide. Peptide-spectrum matches were filtered using up to 2 missed cleavages, unique peptides only, average signal intensity > 5,000 for any pair of technical replicates, and a coefficient of variation < 0.5. Peptides from the different differential-modification searches were pooled and distinctly phosphorylated peptides (e.g., a singly- vs. doubly-phosphorylated version) were kept as separate entries and not averaged. Peptides sharing the same shortest representative sequence for a given S/T/Y residue were averaged (mean).

Channel ratios were first median-normalized across samples within each condition, so that the median signal intensity was equalized across the M0 or TLR4 samples. To isolate changes in phosphorylation from changes in protein abundance, replicate-level phosphosite ratios were then normalized to the whole proteome by dividing the channel ratio from each phosphosite by the corresponding protein’s whole-proteome fold change. Phosphosites lacking whole-proteome values were dropped. Per-site log_2_ fold changes and t-test p-values were computed between conditions. Sites were called significantly phosphorylated at a fold change > 2 (after whole-proteome normalization) and p < 0.05 (independent two-sample t-test).

##### Western blot analysis

Frozen cell pellets were thawed on ice, resuspended in PBS with protease inhibitor cocktail (Roche cOmplete Mini, EDTA-free), and lysed by sonication (3 x 8 pulses, 60% duty cycle). Protein concentration was determined using the DC assay (Bio-Rad), and lysates were normalized to 2 mg/mL. Samples were combined with 4X Laemmli SDS sample buffer (Thermo Scientific, J60015.AD) and boiled at 95 °C for 5 min. Proteins were resolved on a 4-20% gel (Bio-Rad, 4561096 or 5678095) with 1X running buffer (Bio-Rad, 1610732) at 80 V for 20 min, followed by 150 V for 60 min or until the dye front reached the bottom of the gel. Proteins were transferred onto 0.45 µM PVDF membrane (Cytiva) using the Trans-Blot Turbo Transfer System in Transfer buffer (Bio-Rad, 10026938) with methanol in place of ethanol and using the manufacturer’s standard transfer protocol. For total protein staining, membranes were washed with water and dried for 10 min at 37 °C. Membranes were then rehydrated with MeOH before incubation with total protein stain (LI-COR, 926-11011) for 10 min. Membranes were imaged on a ChemiDoc MP Imaging System (Bio-Rad) with the 680 nm channel. Membranes were then blocked for 1 h with 5% milk in TBST (Tris-buffered saline with 0.1% Tween-20). After incubation, membranes were washed once with TBST and incubated with primary antibody for either 2 h at room temperature or overnight at 4 °C. Membranes were then washed with TBST (4 x 5 min, with shaking), then incubated with either goat anti-mouse HRP (Thermo Fisher, G-21040), goat anti-rabbit HRP (Thermo Fisher, G-21234), donkey anti-sheep HRP (Invitrogen, A16041), or mouse anti-rabbit IgG HRP (CST, 3678) for 1 h at room temperature while shaking. Membranes were washed with TBST (4 x 5 min, with shaking), and imaged with substrate (Thermo Scientific, 32106X4 or 34095). Bands were quantified with both Fiji and ImageStudio (LI-COR) and normalized to Vinculin loading control where possible, or to the total protein stain.

###### Modified Western blot for S100A4

For S100A4 blots, the following changes to the standard protocol were made. The proteins were resolved on a 16.5% Criterion Tris-Tricine gel (Bio-Rad, 3450065) at 80 V for 20 min and then at 150 V until the dye front reached the bottom (2-3 h) in Tris-Tricine running buffer. The gel was then transferred to a 0.2 µM nitrocellulose membrane in a wet transfer tank for 60 min at 90 V in the cold room, using a modified transfer buffer (20% Methanol in Tris-Glycine buffer). The membrane was then blocked and stained as described above.

| Marker | Species | Company | Catalog # | Clone | Dilution |
| --- | --- | --- | --- | --- | --- |
| Vinculin | Mouse | Santa Cruz | sc-73614 | 7F9 | 1:200 |
| STAT4 | Rabbit | Cell Signaling Technologies | 2653S | C46B10 | 1:1000 |
| SCO2 | Rabbit | Proteintech | 21223-1-AP | Polyclonal | 1:200 |
| CAMK1 | Sheep | R&D Systems | AF7899 | Polyclonal | 1:200 |
| S100A4 | Sheep | R&D Systems | AF4138 | Polyclonal | 1:200 |
| Tbck (Fy1) | Rabbit | Atlas Antibodies | HPA039951 | Polyclonal | 1:1000 |
| PPP1R21 (Fy2) | Mouse | Schuhmacher et al<sup>6</sup><br>and Quentin et al<sup>7</sup> | -- | -- | 1:1000 |
| C1orf43 (Fy3) | Rabbit | Atlas Antibodies | HPA037871 | Polyclonal | 1:1000 |
| Cryz11 (Fy4) | Rabbit | Schuhmacher et al<sup>6</sup><br>and Quentin et al<sup>7</sup> | -- | -- | 1:7000 |
| DISC1 | Mouse | Santa Cruz | sc-365591 | B-7 | 1:200 |

##### Western blot - ABPP analysis

Frozen cell pellets were thawed on ice, resuspended in PBS, and lysed by sonication (3 x 8 pulses, 60% duty cycle). Protein concentration was determined using a bicinchoninic acid (microBCA) assay (Pierce, 23235), and lysates were normalized to 2 mg/mL. Methyl-PEG-Maleimide Reagent (Thermo Fisher Scientific, 22713; MM(PEG)_24_) was dissolved in DMSO at 250 mM. The metabolites, GTP (Aaron Chemicals, AR00I7U9) and ATP (Aaron Chemicals, AR00I7HK), were dissolved in H_2_O at 125 mM and adjusted to pH 7.5. Each metabolite and H_2_O control (2 µL) were added to 46 µL of macrophage lysates, mixed by pipetting, and incubated for 10 min at room temperature. Following this pre-treatment, 2 µL of maleimide probe or DMSO was added to each sample, mixed by pipetting, and incubated for 30 min at room temperature. Samples were then combined with 4X Laemmli SDS sample buffer (Thermo Scientific, J60015.AD) and boiled at 95°C for 10 min. Proteins were resolved on a 12% gel (Bio-Rad, 4561043) with 1X running buffer (Bio-Rad, 1610732) at 90 V for 15 min, followed by 150 V for 60 min or until the dye front reached the bottom of the gel. Proteins were transferred onto 0.45 µM PVDF membrane (Cytiva) using the Trans-Blot Turbo Transfer System in Transfer buffer (Bio-Rad, 10026938), with methanol in place of ethanol, using the manufacturer’s standard transfer protocol. For total protein staining, membranes were washed with water and dried for 10 min at 37 °C. Membranes were then rehydrated with MeOH before incubation with total protein stain (LI-COR, 926-11011) for 10 min. Membranes were imaged on a ChemiDoc MP Imaging System (Bio-Rad) with the IRDye 680 setting. Membranes were then blocked for 1 h with 5% milk in TBST (Tris-buffered saline with 0.1% Tween-20). After incubation, membranes were washed once with TBST and incubated with primary antibody (Cell Signaling Technology, #2466) for either 2 h at room temperature or overnight at 4 °C. Membranes were then washed with TBST (4 x 5 min, with shaking), then incubated with goat anti-rabbit HRP (Thermo Fisher, G-21234) for 1 h at room temperature while shaking. Membranes were washed with TBST (4 x 5 min, with shaking), and imaged with substrate (Thermo Scientific, 32106X4 or 34095). Bands were quantified with both Fiji and ImageStudio Lite version 5.2 (LI-COR).

##### FERRY Immunoprecipitation for Western blot and mass spectrometry

###### Cell lysis and antibody enrichment

Frozen cell pellets were thawed on ice, resuspended in lysis buffer (50 mM Tris pH 7.4, 150 mM NaCl, 1% Triton X-100), and incubated at 4 °C with rotation. Tubes were then centrifuged to remove debris (16,000 x *g*, 4 min, 4 °C), and the supernatant was transferred to another tube. Protein concentration was determined by DC assay, and lysates were normalized to 2 mg/mL with lysis buffer. Lysate (500 µL, 1 mg) was transferred into separate low-binding Eppendorf tubes for each treatment condition. Primary antibodies (Fy2 or Fy1 (Atlas Antibodies, HPA039951); 1.5 µg/tube) or Normal IgG (CST, Rabbit (2729) or Mouse (68860)) were then added to the lysate, and the samples were incubated at 4 °C for 3 h with rotation.

###### Dynabead incubation and bead washes

After 3 h, 20 µL of washed Dynabeads Protein A (for rabbit antibodies, Invitrogen, 10002D) or Protein G (for mouse antibodies, Invitrogen, 10003D) were added per sample, and the samples were rotated for another 45 min. Tubes were spun briefly (500 x *g*, 5 s, 25 °C) to remove any remaining liquid from the lids and transferred to a magnetic rack. The flow-through was saved to validate target protein depletion. Beads were then washed 4 times with wash buffer (50 mM Tris pH 7.4, 150 mM NaCl, 0.2% Triton X-100). Samples prepared for Western blot analysis were then incubated with 1X Laemmli SDS sample buffer diluted in PBS for 10 min at 95 °C. Samples were spun briefly (500 x *g*, 5 s, 25 °C), placed on the magnet, and the supernatant was transferred to a new low-bind Eppendorf tube and stored at 4 °C until further analysis.

Samples prepared for downstream mass spectrometry analysis were washed an additional two times with 200 mM EPPS (pH 8.0). Samples were then eluted with 60 µL of 8 M urea in 200 mM EPPS (pH 8.0) and heated at 65 °C for 10 min. Samples were placed on the magnet, and the supernatant was transferred to a new low-bind Eppendorf tube. Freshly made DTT (3.75 µL, 200 mM in water) was added to each sample and incubated for 15 min at 65 °C. Freshly made iodoacetamide (3.75 µL, 400 mM in water) was then added and the samples were incubated at 37 °C in the shaker for 30 min. Samples were then diluted with 202.5 µL of EPPS (pH 8.0) and digested with 4 µg of Trypsin/LysC and 2 µL of 100 mM CaCl_2_ overnight with shaking.

###### TMT labeling

Acetonitrile (90 µL) was added to digested samples for a final volume of 25%. TMT reagents (5 µL, 20 µg/µL) were then added to each tube and incubated for 1 h at room temperature. The reaction was quenched with 5 µL of 5% hydroxylamine for 15 min at RT, then acidified with 12 µL of formic acid. Samples were combined, split back into 2 tubes, and dried in the SpeedVac. Samples were desalted using C18 SepPak cartridges and fractionated as described above. Following fractionation, samples were combined into six final fractions and analyzed by LC-MS/MS/MS as described above.

###### Data processing and analysis

To correct for differences in pull-down efficiency between samples, signal intensities in each IP (TBCK or PPP1R21) were scaled by a per-donor normalization factor derived from the median signal of the bait protein, so that bait levels were equalized across conditions. Percent control values were calculated as a ratio to the per-donor median of the corresponding M0-IP control. Per-protein fold changes were computed as the log_2_-transformed ratios of mean intensities between the two conditions, and significance was assessed by an independent two-sample t-test. Proteins were called enriched when FC > 1.5 and p < 0.05.

Background contaminants were identified using the CRAPome (reprint-apms.org, v2.0)^8^, querying all quantified proteins against the 716 human control experiments; a protein detected in more than 200 of these was flagged as a CRAPome hit. Enriched protein sets were restricted to non-CRAPome proteins prior to downstream GO-term enrichment.

##### Immunofluorescence

###### Coverslip Coating

12 mm round glass coverslips (#1.5, Corning) were sterilized in 70% Ethanol and transferred to each well of a 24-well plate. Coverslips were washed twice with molecular biology grade water and then incubated with 0.5 mL of Poly-*L*-Lysine (Sigma-Aldrich, 50-176-1647) resuspended at 0.1 mg/mL in molecular biology grade water for 20 min at room temperature. Coverslips were then washed twice in molecular biology grade water and transferred to a new 24-well plate to dry. After coverslips were fully dried, they were used immediately for cell plating.

###### Macrophage plating and polarization

Macrophages that were polarized for 6 days, washed once with PBS, and incubated with TrypLE for 30 min in a humidified 37 °C incubator. After 30 min, cells were gently scraped, transferred into a Falcon tube, and centrifuged (520 x *g*, 5 min, 25 °C). Cells were resuspended in PBS, counted, centrifuged again (520 x *g*, 5 min, 25 °C), and resuspended at 150,000 cells / mL in macrophage media containing mCSF. Cells (0.5 mL, 75,000 cells per well) were then transferred to a 24-well plate with pre-coated Poly-*L*-Lysine glass coverslips and left to adhere overnight at 37 °C in a humidified incubator. The next day, media was aspirated from the plates, and cells were re-supplemented with either macrophage media with mCSF or media with mCSF and LPS at 100 ng/mL and returned to the incubator for 24 h. Coverslips containing adherent cells were then washed twice with PBS-MC (PBS with 1 mM MgCl_2_ and 0.1 mM CaCl_2_) before incubation with freshly made 4% PFA (pH 7.0) in PBS-MC for 15 min at room temperature. Coverslips were then washed again with PBS-MC and kept at 4 °C until staining.

###### Immunofluorescence staining

Cells were permeabilized with 0.1% Triton X-100 in PBS-MC for 5 min at room temperature. Coverslips were then washed once with PBS-MC and blocked with 2% Normal goat serum (NGS, Jackson Immunology, 005-000-121) and 2% BSA in PBS-MC for 1 h at room temperature. Coverslips were then washed twice with PBS-MC and transferred to a humidified chamber with the adherent cells facing up. Staining solutions in blocking buffer supplemented with 0.1% Triton X-100 (60 µL) were gently layered onto the surface of each coverslip with a pipette, and the chamber was incubated overnight at 4 °C. The next day, the coverslips were transferred back into the 24-well plate and washed twice with PBS-MC. Cells were then incubated with fluorescently labeled secondary antibodies in blocking buffer with 0.1% Triton X-100 for 1 h at room temperature in the dark. Coverslips were then washed twice with PBS-MC and incubated with DAPI (Invitrogen R37606, 2 drops per mL of blocking buffer) for 15 min at room temperature. Coverslips were then washed five times with PBS-MC and mounted onto glass slides with ProLong Diamond Antifade Mountant (Invitrogen, P36961). Sealed coverslips were then imaged on a Zeiss LSM980 confocal microscope with Airyscan 2 detector. All images reported here were acquired in conventional confocal mode with GaAsP-PMT (green/red/far-red dyes) or multialkali PMT-based detection for DAPI. A Plan-Apochromat 63x/1.40 NA oil-immersion objective was used with immersion oil of refractive index 1.518. Alexa Fluor 488 and Alexa Fluor 568 were excited at 488 nm and 561 nm and detected over spectral bandwidth of 498-552nm, and 568-640nm respectively; DAPI was excited at 405 nm and detected over 410-482nm. The confocal pinhole was set to 64.9 µm. Images were acquired at 16-bit, with bidirectional scanning, a frame size of 1908x1908 pixels and an optical zoom of 1X. Z-stacks of 15-40 planes were acquired to span the entire cell height. Channels were acquired sequentially to prevent spectral bleed-through. Laser power, detector gain, digital offset, pinhole diameter, scan speed, line averaging and optical zoom were kept identical across conditions.

| Marker | Antibody Species | Catalog # | Dilution | Secondary Fluorophore |
| --- | --- | --- | --- | --- |
| Fy1 | Rabbit | HPA039951<br>(Atlas Antibodies) | 1:200 | Goat anti-Rabbit AF568 |
| Fy2 | Mouse | Schuhmacher et al<sup>6</sup><br>and Quentin et al<sup>7</sup> | 1:1000 | Goat anti-Mouse AF488 |
| Fy4 | Rabbit | Schuhmacher et al<sup>6</sup><br>and Quentin et al<sup>7</sup> | 1:7000 | Goat anti-Rabbit AF568 |
| Rab5 | Rabbit | 3547 (CST) | 1:200 | Goat anti-Rabbit AF568 |

###### IF analysis

First, 3D image stacks were denoised by N2V neural network implemented in MotionTracking (https://motiontracking.mpi-cbg.de), which improved Signal-to-Noise Ratio (SNR) on ∼30dB. Then 3D image stacks were deconvolved plane by plane by DeconSVD (MotionTracking) with theoretical microscope point-spread function (NA=1.4 and emission wavelength was taken from microscope metadata). Then background was subtracted^9^ and fluorescent puncta were segmented as 3D objects in MotionTracking v.8.102.3. The segmentation threshold was set at 15% of detected vesicle intensity level.

Whole-cell masks were defined for the Fy2 channel by thresholding at an intensity of 1000. The number of puncta and average puncta volume were determined per cell. For endocytosis analysis, puncta with < 0.15 µm radius were filtered out and Rab5 puncta volume and number of Rab5 puncta per cell volume were calculated per cell, with at least 25 cells quantified per donor and condition across six donors. Individual cells were plotted on a superplot,^10^ with the average of all cells reported per donor. Significance was calculated by a two-tailed paired t-test on the donor means, and average and SEM of six donors was plotted.

For colocalization, puncta with < 0.15 µm radius were filtered out, and colocalization was calculated as fraction of integral intensity associated with colocalized objects^11^ with a 25% apparent object volume overlap cutoff. Given that deconvolution was performed in 2D (plane-by-plane) and Z-dimension was not deconvolved, the “radius” was defined as 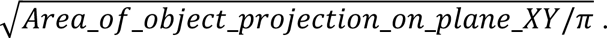

Colocalization was determined separately for both markers (i.e., Rab5 overlap with Fy2 and Fy2 overlap with Rab5) on a per-cell basis, with at least 25 cells quantified per donor and condition across six donors. Individual cells were plotted on a superplot, with the average of all cells reported per donor. Significance was calculated by a two-tailed paired t-test on the donor means, and average and SEM of six donors was plotted. Per-cell measurements and donor-level statistics are provided in Data S4.

##### Proximity Ligation Assay

Day 6 macrophages were lifted as described above and plated on a 96-well glass bottom plate (Cellvis, P961.5HN) at 15,000 cells per well in macrophage media with mCSF and left to adhere overnight in a humidified incubator chamber at 37 °C. Media was then removed from the wells, and cells were polarized with either macrophage media with mCSF alone, or mCSF with LPS at 100 ng/mL. After 24 h incubation, the plate was washed twice with room temperature PBS-MC, and cells were fixed with 4% PFA in PBS-MC for 15 min at room temperature. Cells were then washed twice with PBS-MC and stored at 4 °C until staining.

###### Membrane labeling, permeabilization, and blocking

Cells were labeled with 5 µg/mL wheat germ agglutinin (WGA-AF488, Invitrogen W11261) in PBS-MC for 15 min at room temperature, washed twice with PBS-MC, and permeabilized with 0.1% Triton X-100 in PBS-MC for 5 min at room temperature. Cells were washed twice more and blocked with Duolink Blocking Solution (Sigma-Aldrich, DUO82007) for 1 h at 37 °C. Blocking solution was then removed, and primary antibodies diluted in Duolink Antibody Diluent (Sigma-Aldrich, DUO82008) were added and incubated overnight at 4 °C.

###### PLA probe incubation, ligation, and amplification

The next day, cells were washed twice for 5 min each in Wash Buffer A (0.01 M Tris, 0.15 M NaCl, 0.05% Tween-20). Duolink Probe Anti-Rabbit PLUS (DUO92002-100RXN) and Probe Anti-Mouse MINUS (DUO92004-100RXN) were diluted at a 1:5 ratio in Duolink Antibody Diluent and incubated for 20 min at room temperature before addition to cells (40 µL per well). Plates were incubated in a humidified chamber for 1 h at 37 °C. Cells were then washed twice with Wash Buffer A, and 40 µL of Ligation mix (ligase diluted 1:40 in 1X Ligation buffer; DUO92008-100RXN) was added to each well. The plate was incubated in a pre-heated humidity chamber for 30 min at 37 °C. Cells were then washed twice with Wash Buffer A, and 40 µL of Amplification mix (polymerase diluted 1:80 in 1X Amplification buffer) was added to each well, followed by an additional incubation in a pre-heated humidified chamber for 100 min at 37 °C.

###### Washing and mounting

Cells were washed twice with Wash Buffer B (0.2 M Tris, 0.1 M NaCl) for 10 min at room temperature, then washed once with 0.01X Wash Buffer B for 1 min. Cells were then mounted with ProLong Diamond Antifade Mountant with DAPI (Invitrogen, P36962).

###### PLA analysis

PLA samples in 96-well glass-bottom plates were imaged on the same LSM980 microscope used for immunofluorescence with acquisition settings held identical across all panels, conditions, donors and sessions; acquisition parameters were otherwise as described for immunofluorescence above. Images were deconvolved in SVI Huygens Professional (Scientific Volume Imaging, V26.04) using the Classic Maximum Likelihood Estimation (CMLE) algorithm with a theoretical point-spread function computed from the microscopic parameters discussed above, voxel dimensions, and embedding medium refractive index (approx. 1.47). These parameters were used by the software to deduce signal-to-noise ratio and a quality threshold stopping criteria for deconvolution algorithm. Deconvolution iterations were run until the quality criterion was fulfilled. Identical deconvolution parameters were applied to all conditions and all donors.

| Panel | Marker | Antibody Species | Catalog # | Dilution |
| --- | --- | --- | --- | --- |
| 1 | Fy1 | Rabbit | HPA039951 (Atlas Antibodies) | 1:200 |
| 1 | Fy2 | Mouse | Schuhmacher et al<sup>6</sup> and Quentin et al<sup>7</sup> | 1:1000 |
| **2** | **Fy3** | **Rabbit** | **APrEST80927 (Atlas Antibodies)** | **1:200** |
| **2** | **Fy2** | **Mouse** | Schuhmacher et al<sup>6</sup> and Quentin et al<sup>7</sup> | **1:1000** |
| 3 | Fy4 | Rabbit | Schuhmacher et al<sup>6</sup> and Quentin et al<sup>7</sup> | 1:7000 |
| 3 | Fy2 | Mouse | Schuhmacher et al<sup>6</sup> and Quentin et al<sup>7</sup> | 1:1000 |
| **4** | **Rab5** | **Rabbit** | **3547 (CST)** | **1:200** |
| **4** | **Fy2** | **Mouse** | Schuhmacher et al<sup>6</sup> and Quentin et al<sup>7</sup> | **1:1000** |
| 5 | DISC1 | Rabbit | 40-6900 (Invitrogen) | 1:200 |
| 5 | Fy2 | Mouse | Schuhmacher et al<sup>6</sup> and Quentin et al<sup>7</sup> | 1:1000 |

##### Generation of reference protein tables

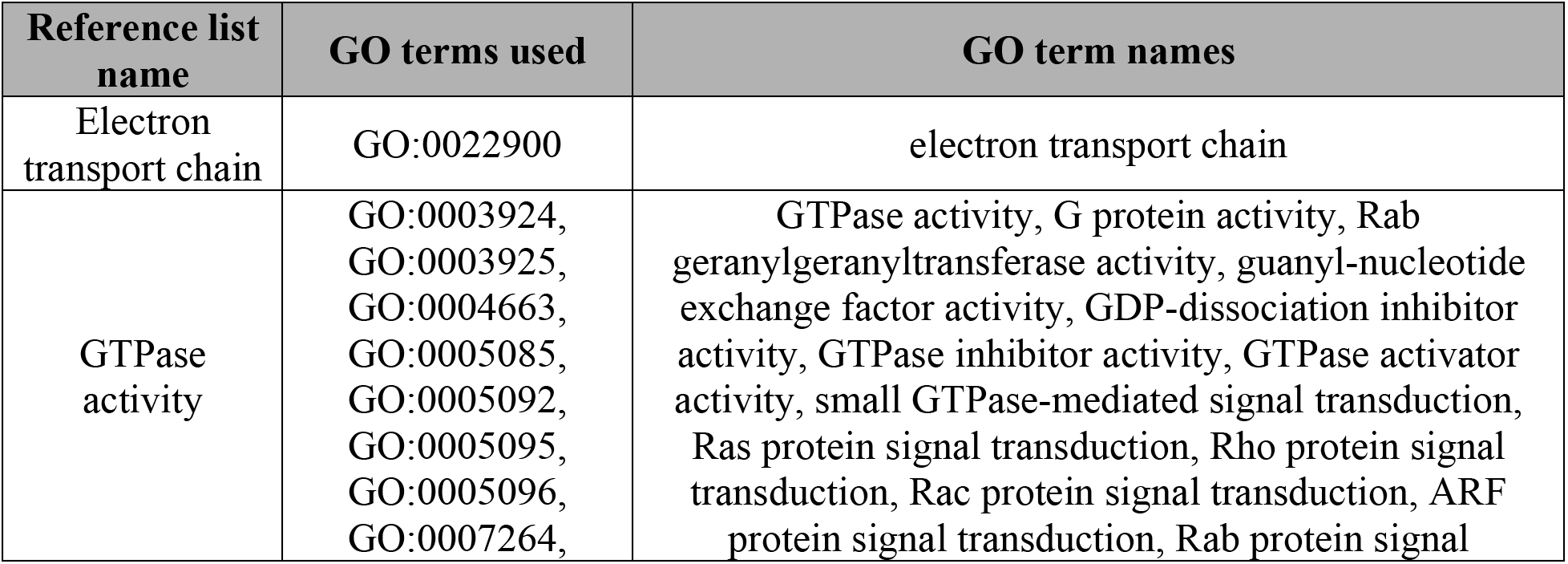
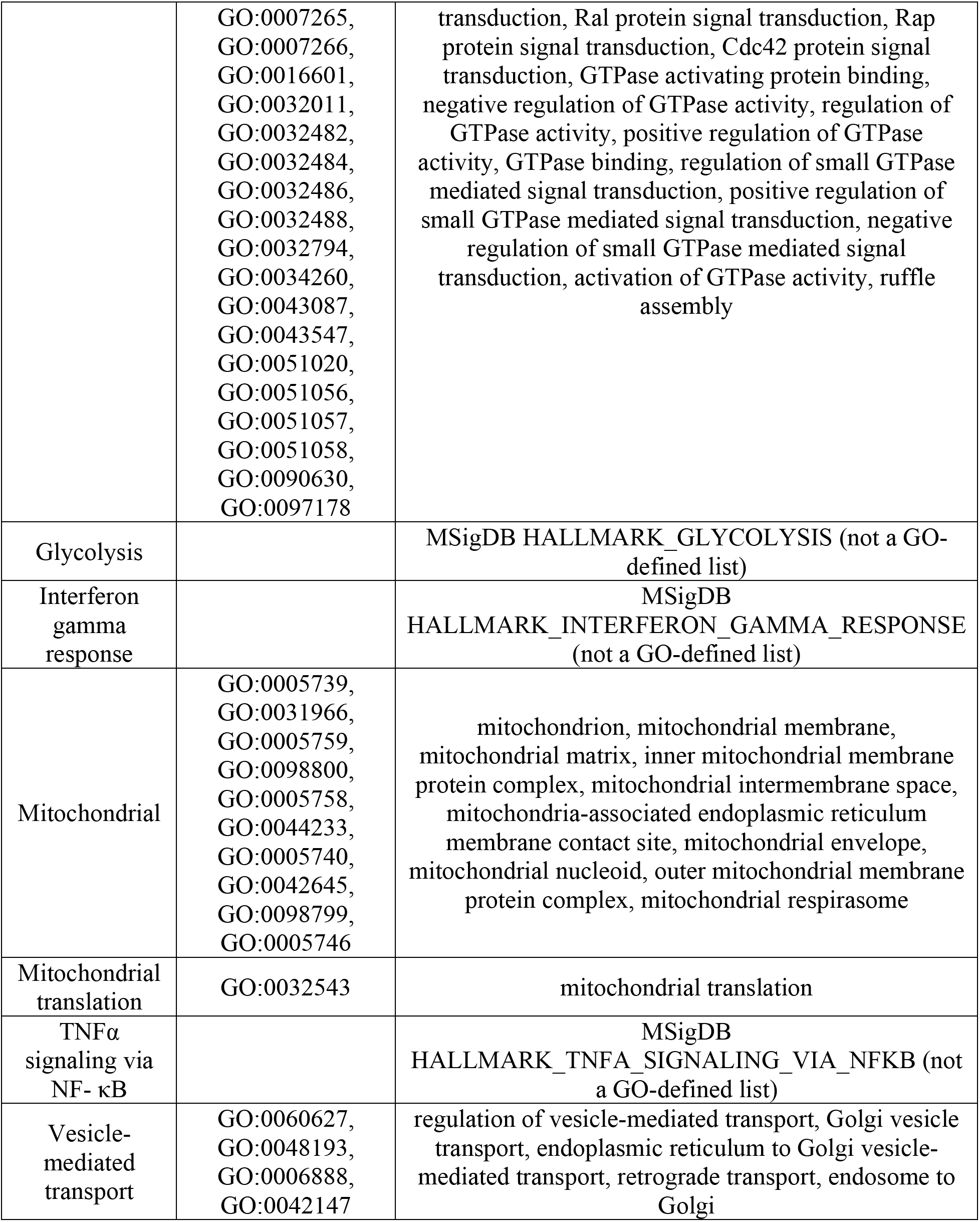

### (C) Key resources

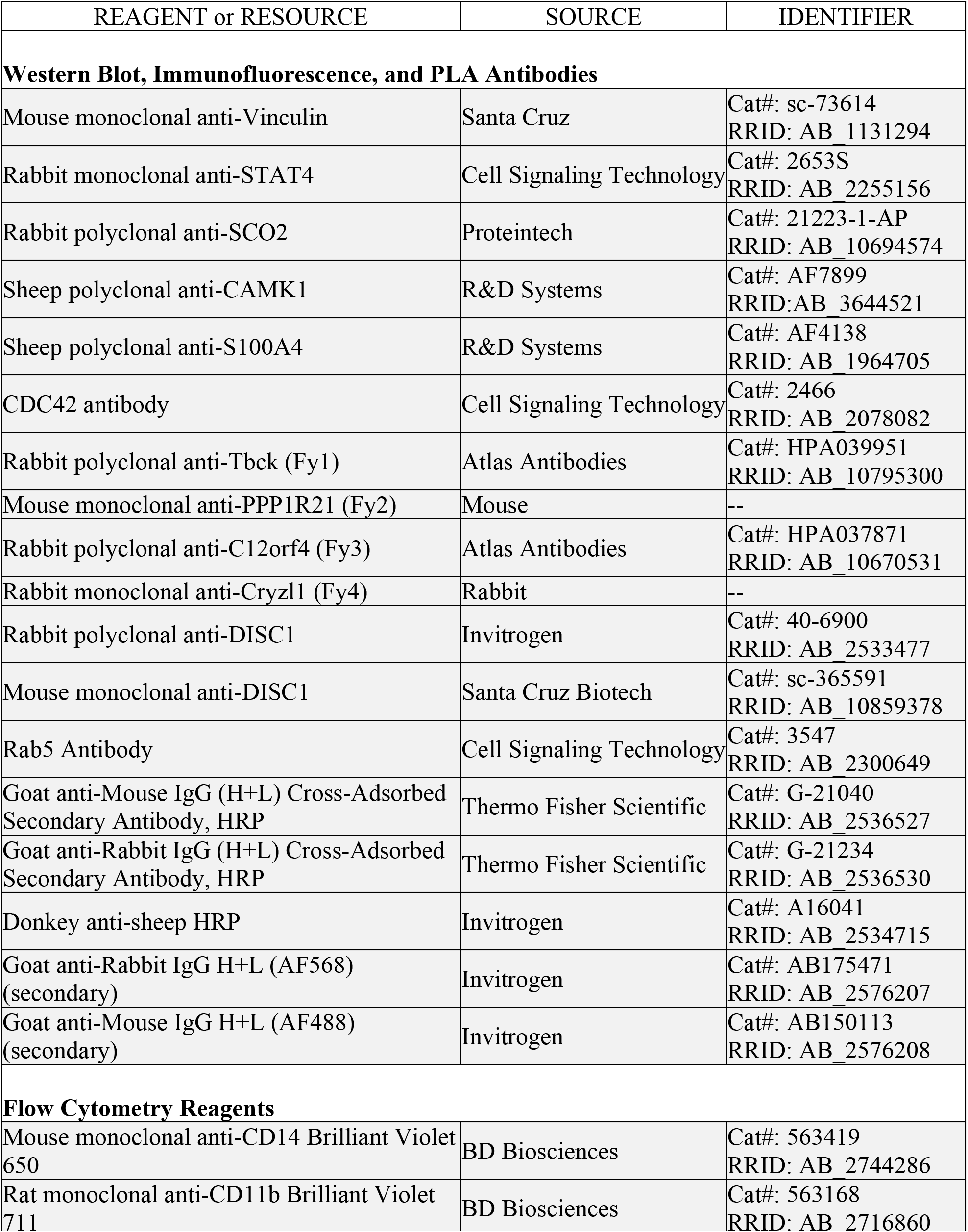
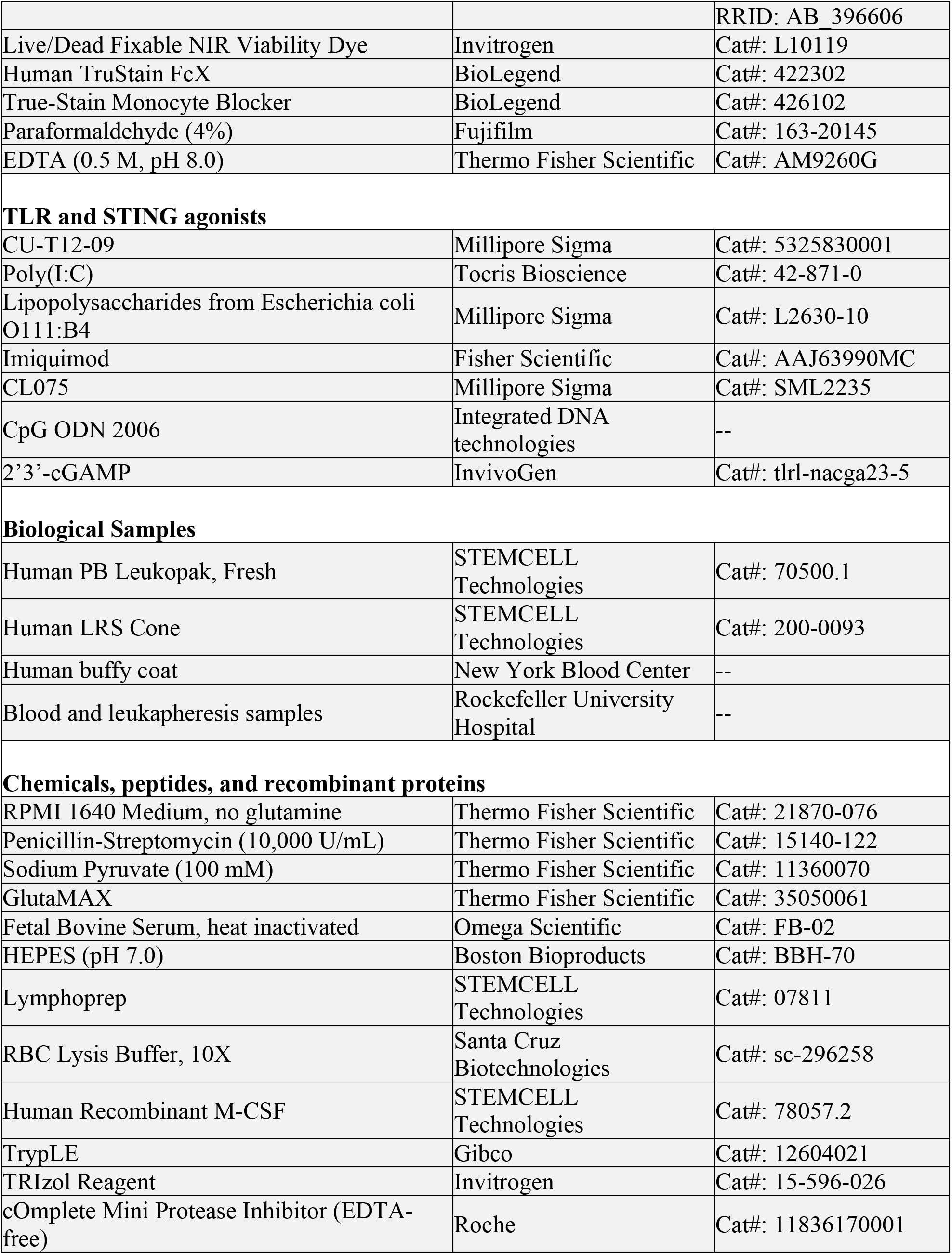
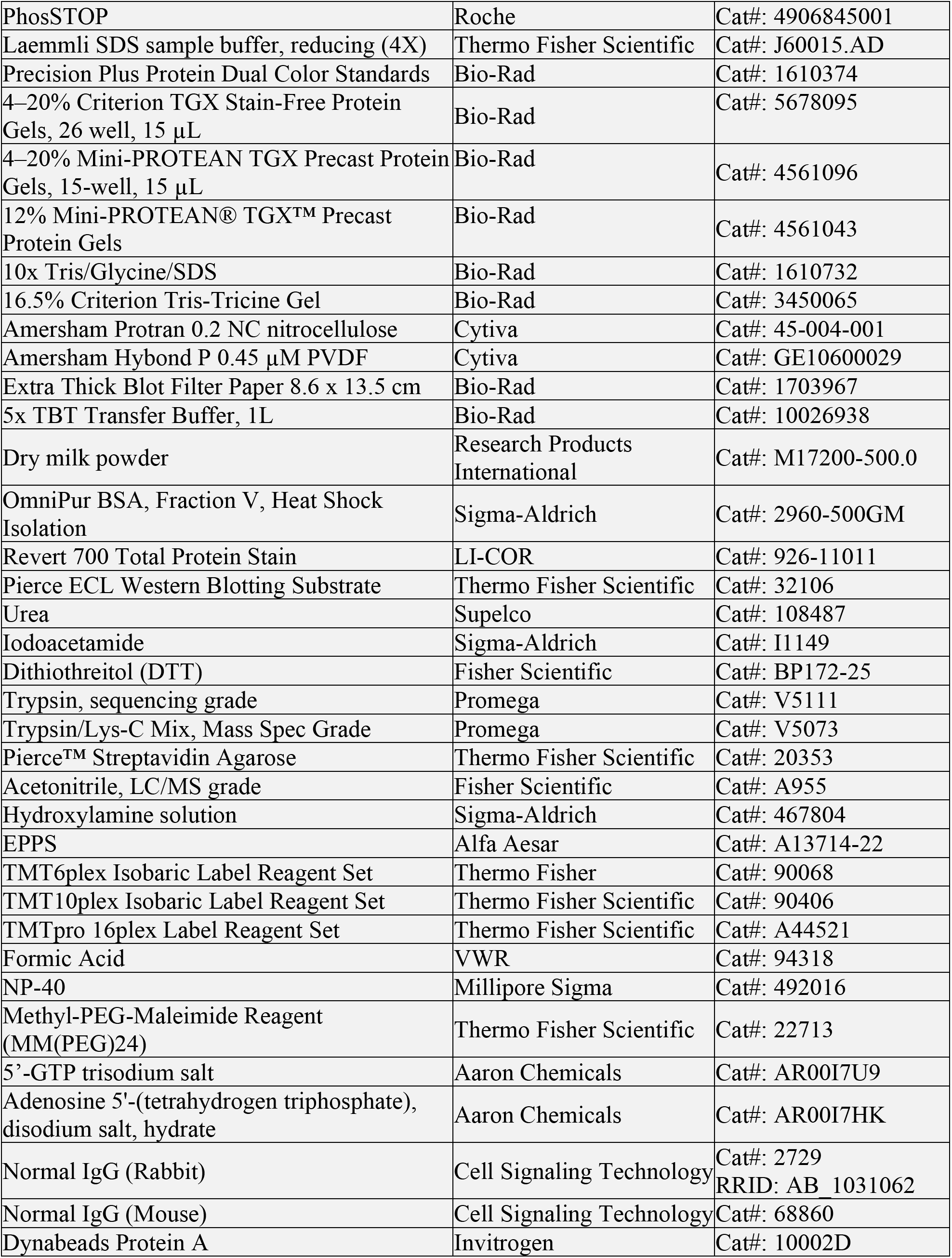
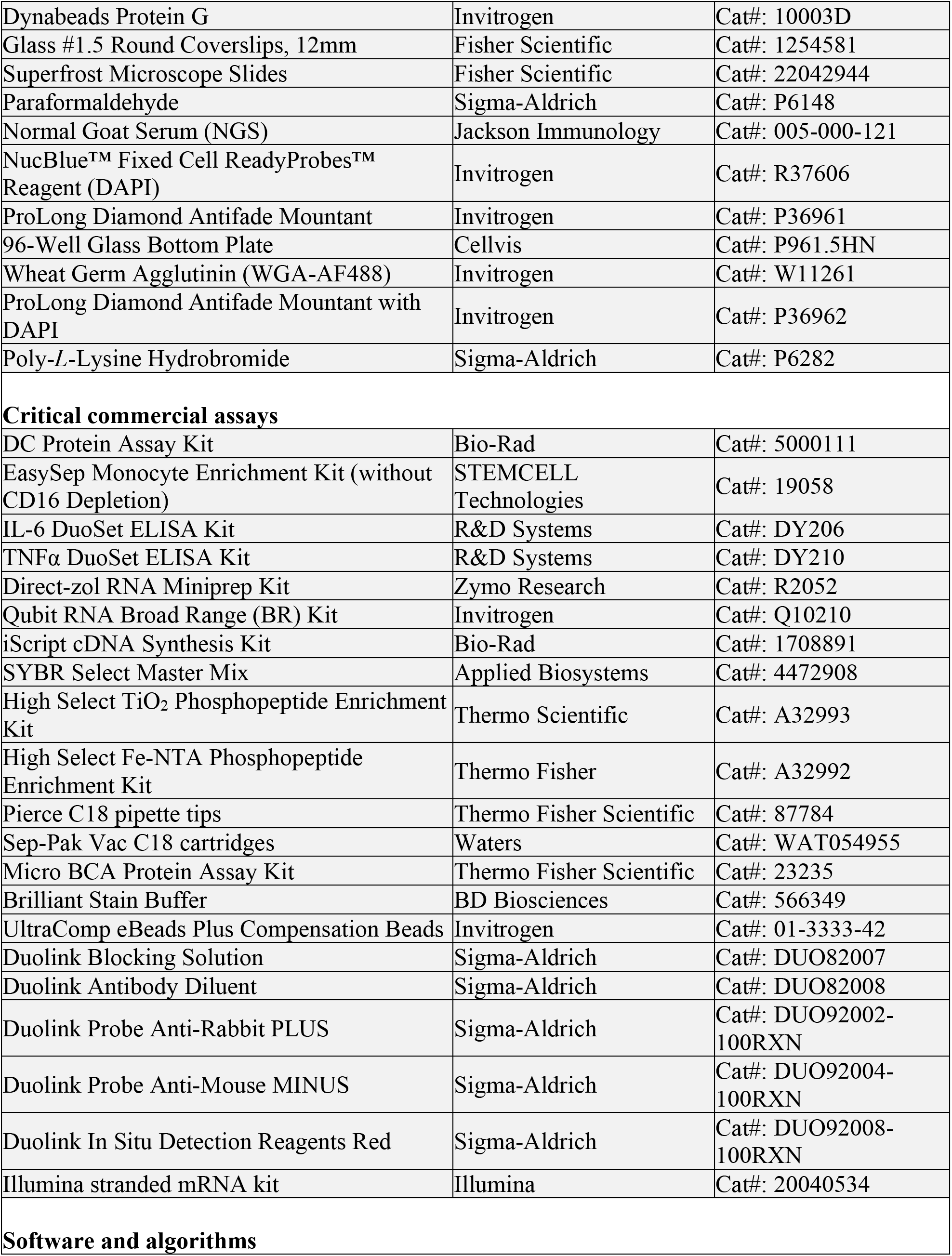
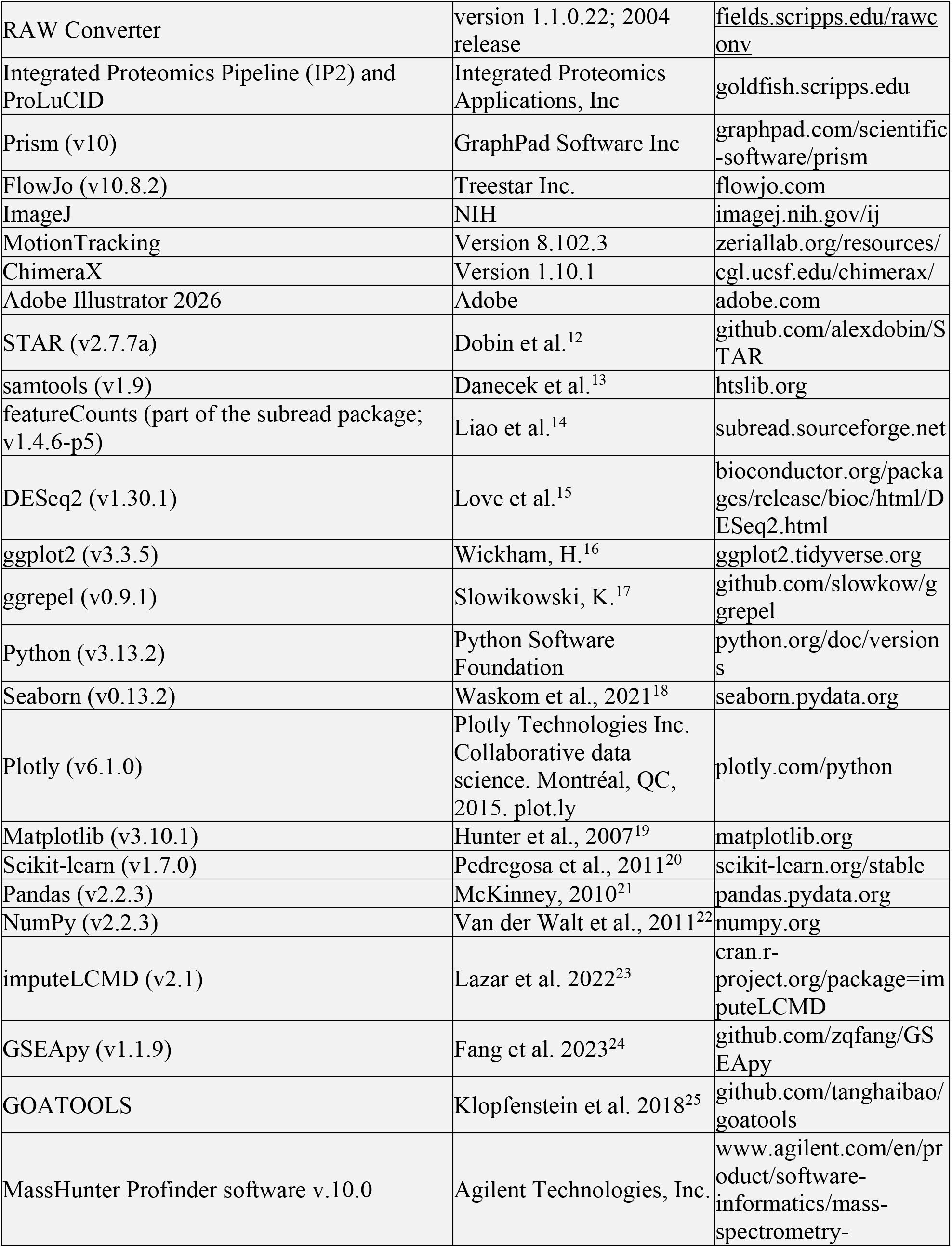
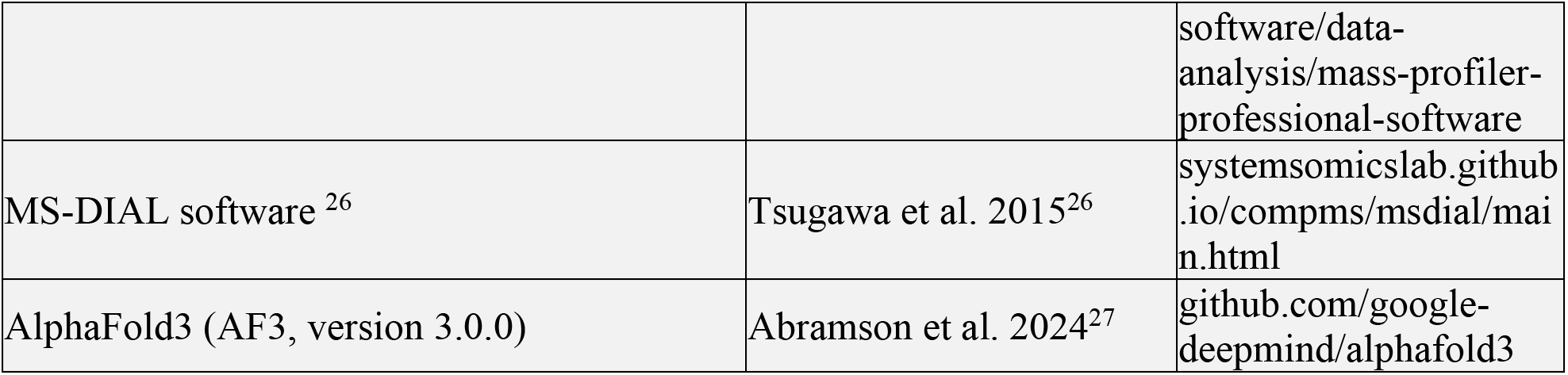

